# Genetic control of variability in subcortical and intracranial volumes

**DOI:** 10.1101/443549

**Authors:** Aldo Córdova-Palomera, Dennis van der Meer, Tobias Kaufmann, Francesco Bettella, Yunpeng Wang, Dag Alnæs, Nhat Trung Doan, Ingrid Agartz, Alessandro Bertolino, Jan K. Buitelaar, David Coynel, Srdjan Djurovic, Erlend S. Dørum, Thomas Espeseth, Leonardo Fazio, Barbara Franke, Oleksandr Frei, Asta Håberg, Stephanie Le Hellard, Erik G. Jönsson, Knut K. Kolskår, Martina J. Lund, Torgeir Moberget, Jan E. Nordvik, Lars Nyberg, Andreas Papassotiropoulos, Giulio Pergola, Dominique de Quervain, Antonio Rampino, Genevieve Richard, Jaroslav Rokicki, Anne-Marthe Sanders, Emanuel Schwarz, Olav B. Smeland, Vidar M. Steen, Jostein Starrfelt, Ida E. Sønderby, Kristine M. Ulrichsen, Ole A. Andreassen, Lars T. Westlye

## Abstract

Sensitivity to external demands is essential for adaptation to dynamic environments, but comes at the cost of increased risk of adverse outcomes when facing poor environmental conditions. Here, we apply a novel methodology to perform genome-wide association analysis of mean and variance in nine key brain features (accumbens, amygdala, caudate, hippocampus, pallidum, putamen, thalamus, intracranial volume and cortical thickness), integrating genetic and neuroanatomical data from a large lifespan sample (n=25,575 individuals; 8 to 89 years, mean age 51.9 years). We identify genetic loci associated with phenotypic variability in cortical thickness, thalamus, pallidum, and intracranial volumes. The variance-controlling loci included genes with a documented role in brain and mental health and were not associated with the mean anatomical volumes. This proof-of-principle of the hypothesis of a genetic regulation of brain volume variability contributes to establishing the genetic basis of phenotypic variance (i.e., heritability), allows identifying different degrees of brain robustness across individuals, and opens new research avenues in the search for mechanisms controlling brain and mental health.

## Introduction

Phenotypic variability is key in evolution, and partly reflects inter-individual differences in sensitivity to the environment ^1^. Genetic studies of human neuroanatomy have identified shifts in mean phenotype distributions (e.g., mean brain volumes) between groups of individuals with different genotypes ^2^, and have documented genetic overlaps with common brain and mental disorders ^3^. Despite the evolutionary relevance of phenotypic dispersion evidenced in multiple species and traits ^1, 4^, the genetic architecture of variability in human brain morphology is elusive.

Phenotypic variance across genotypes can be interpreted in relation to robustness, i.e., the persistence of a system under perturbations ^1, 4^ and evolvability, the capacity for adaptive evolution ^5^. High phenotypic robustness is indicated by low variation in face of perturbations, i.e. phenotypes are strongly determined by a given genotype. In contrast, lack of robustness corresponds to high sensitivity, yielding phenotypes with overall larger deviations from the population mean in response to environmental, genetic or stochastic developmental factors. Neither increased or decreased robustness confers evolutionary advantages *per se* ^1^, and their consequences for adaptation need to be understood in view of the genotype-environment congruence. Reduced robustness (and thus increased variability of trait expression) can be a conducive to adaptive change ^5^, and increased variability of phenotypic expression can in itself also be favored by natural selection in fluctuating environments ^6^. Thus, recognizing genetic markers of sensitivity can aid in identifying individuals who are more susceptible to show negative outcomes when exposed to adverse factors –either genetic or environmental– and otherwise optimal outcomes in the presence of favorable factors. Such variance-controlling genotypes may be conceived as genomic hotspots for gene-environment and/or gene-gene interactions, with high relevance for future genetic epidemiology studies ^7^.

To provide a proof-of-principle of the hypothesis of a genetic regulation of brain volume variability, we conducted a genome-wide association study of intragenotypic variability in seven key subcortical regions and intracranial volume (ICV) using a harmonized genotype and imaging data analysis protocol in a lifespan sample (n=25,575 individuals; 8 to 89 years, mean age 51.9 years; 48% male, Methods and Supplementary Information).

## Materials and Methods

### Participants

Data from 25,575 unrelated European-ancestry individuals were included (mean age 51.9 years, ranging from 8 to 89 years old; 48% male), recruited through 16 independent cohorts with available genome-wide genotyping and T1-weighted structural MRI. Extended information on each cohort reported in Supplementary Information includes recruitment center, genotyping and brain imaging data collection, sample-specific demographics, distribution of brain volumes and, when relevant, diagnoses (795 individuals had a diagnosis). Written informed consent was provided by the participants at each recruitment center, and the protocols were approved by the corresponding Institutional Review Boards.

### Genotypes

Only participants with European ancestry (as determined by multidimensional scaling) were included in the final set of analyses, in recognition that the inclusion of subjects from other ethnicities can potentially add genetic and phenotypic confounding. Except for the UK Biobank cohort, all directly genotyped data were imputed in-house using standard methods with the 1000 Genomes European reference panel. After imputation, each genotyping batch underwent a quality control stage (MAF < 0.01; Hardy-Weinberg equilibrium *p* < 10^−6^; INFO score < 0.8). When all samples were combined, over 5 million distinct markers passed quality control genome-wide. Additional filters on genotyping frequencies were applied to the final merged dataset based on statistical considerations for genotype frequency in variance-controlling detection, as described below. Genetic data analysis was conducted using PLINK ^8^, with R ^9^ plugin functions when appropriate (https://www.cog-genomics.org/plink/1.9/rserve).

### Brain features

Three-dimensional T1-weighted brain scans were processed using FreeSurfer ^10^ (v5.3.0; http://surfer.nmr.mgh.harvard.edu). Mean cortical thickness and eight well-studied volumetric features were selected for analysis moving forward, as literature findings on large datasets show that their mean population value is influenced by common genetic variation ^2^: accumbens, amygdala, caudate, hippocampus, pallidum, putamen, thalamus and ICV. Cohort-wise distribution of values is summarized in Supplementary Information. Before the ensuing statistical analyses, outliers (+-3 standard deviations from the mean) were removed, and generalized additive models (GAM) were implemented in R (https://www.r-project.org) to regress out the effects of scanning site, sex, age, diagnosis and ICV (for subcortical volumes only). Hereafter, brain volumes correspond to residuals from those GAM fits unless otherwise specified.

### Statistical analyses

Genome-wide association statistics were computed for genetic effects on the mean and variance of the volumetric feature distributions. For each marker, the distribution of each outcome phenotype was normalized via rank-based inverse normal transformation (INT) to prevent statistical artifacts. Scale transformations like INT have been shown to aid genetic discovery by constraining mean-effects and reducing the effect of phenotypic outliers, which reduces Type I error rates without sacrificing power ^11, 12^. In short, INT was applied to transform each subject’s phenotype (*y*_*i*_) as

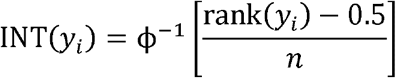

as where rank(*y*_*i*_) is the rank within the distribution, *n* stands for sample size (without missing values) and ϕ^−1^ denotes the standard normal quantile function. Intuitively, all phenotype values are ranked and the ranks are mapped to percentiles of a normal distribution. Then, an additive genetic model was computed with

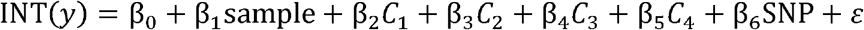

where INT(*y*) is the normalized phenotype variable; SNP is the relevant marker coded additively and ε stands for regression residuals. Four genomic principal components (*C*_1_-*C*_4_) were included, to control for population stratification and cryptic relatedness, and to make the results consistent/comparable with a previous large-scale analysis of genetic variation and brain volumes ^2^. Results from that analysis (mean-model) were contrasted with the statistics from the variance-model. The previous residuals ε were again inverse normal transformed, and used as input for the variance-model using the Brown–Forsythe test. Briefly, INT-transformed residuals were used to compute 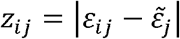, with 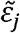 as the median of group *j* (here, genotype) and these, in turn, to compute the *F* statistic:

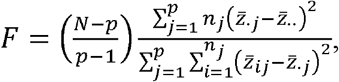

where *n*_*j*_ is the number of observations in group *j*, *p* is the number of groups (2 or 3 different genotypes), and 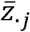 denotes the mean in group *j*. To prevent increases in false positive rates arising from small groups ^13^, only markers with at minimum (non-zero) genotype count of at least 100 were included. This value was chosen based on literature about power and statistical considerations of genome-wide association studies for phenotypic variability ^13^. The data were analyzed and visualized in R with the aid of appropriate packages. When relevant, significant markers were annotated and additionally inspected using FUMA ^14^.

## Results

Genome-wide association statistics were computed for genetic effects on the variance and mean of the volumetric feature distributions. Consistent with previous large-scale analyses on genetics of neuroimaging volumetric measures ^2, 15^, features included bilateral (sum of left and right) amygdala, caudate nucleus, hippocampus, nucleus accumbens, pallidum, putamen and thalamus, as well as ICV and mean cortical thickness. 96.9% of the included participants were healthy controls (n=24,780); the remaining 3.1% were diagnosed with a brain disorder (n=795; including psychosis, depression, and attention deficit hyperactivity disorder, Supplementary Information). The analyses were conducted in a two-stage protocol. For each genotype, we conducted a standard association test for the inverse-normal transformed (INT) brain volumes ^11^, adjusting for scanning site, sex, age, age squared, diagnosis, and ICV (for the subcortical volumes only). Residuals from that model were then INT-transformed and submitted to genome-wide Levene’s tests to investigate if specific alleles associate with elevated or reduced levels of phenotypic variability. For relevant markers, variances explained by mean and variance models were estimated from the INT-transformed volumes before fitting regression models using a previously reported approach ^7^.

A mega-analysis of 25,575 unrelated subjects of European ancestry identified candidate loci associated with differential levels of phenotypic variability overall on four out of the nine volumetric features (pallidum, mean thickness, ICV and thalamus), including two at genome-wide significance (*p*<5×10^−8^), and two with marginal significance (*p*<7 × 10^−8^) (Figure 1). Genomic inflation factors (lambda) ranged between 1.009 and 1.049 for the nine different variance-GWAS (Supplementary Information). A conventional mean phenotype GWAS with additive model on the same set of variants, with INT-transformed phenotypes, showed 56 significant loci influencing four volumetric traits after adjustment for genomic inflation (accumbens [2], amygdala [4], caudate [9], hippocampus [10], pallidum [5], putamen [6], thalamus [2], ICV [10] and mean cortical thickness [8]) (Supplementary Information). Manhattan plots for both mean- and variance-GWAS are displayed as Supplementary Information.

**Figure 1.**
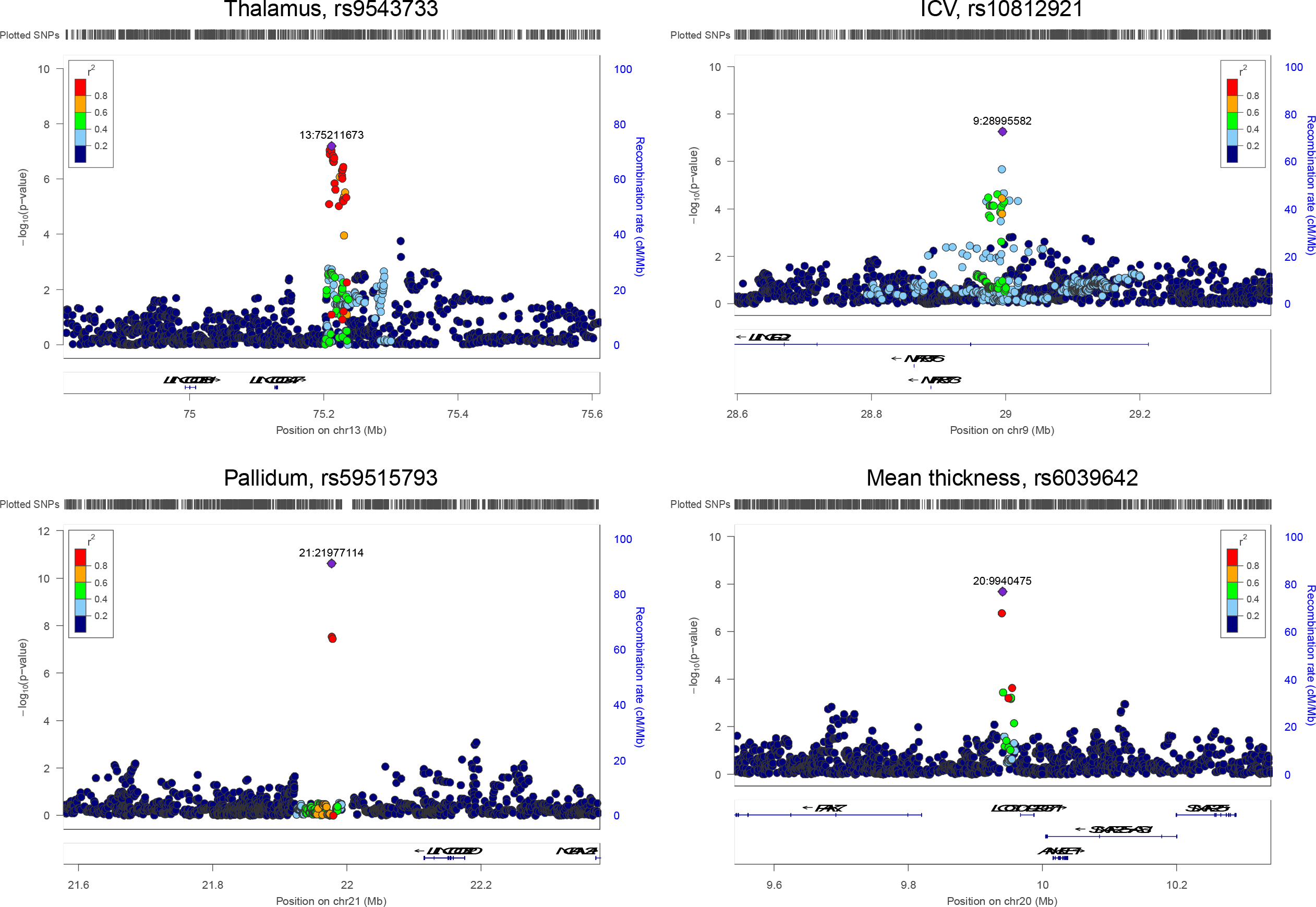
Common genetic variants regulate the distribution variance of human subcortical and intracranial volumes.

In the variance analysis, the top loci included an intergenic region on 21q21.1 around rs59515793 associated with pallidum volume variance (chr21:21977114:G:A; minor allele frequency (MAF)=0.42; 393518 bp from *NCAM2.6*; *p*=2.4×10^−11^; variance explained variance model: 0.405%; variance explained mean model: 0.0005%), and a locus in chromosome 20 between *SNAP25*, *PAK7* and *ANKEF1* associated with mean cortical thickness variability (rs6039642; chr20:9940475:G:A; MAF=0.17; *p*=2.1×10^−8^; variance explained variance model: 0.259%; variance explained mean model: 0.005%). In addition, two loci showed borderline significance associations with variance in neuroanatomical phenotypes: thalamus variability was related to genotypes on an intergenic locus near *LINC00347* (rs9543733; chr13: 75211673:C:T; MAF=0.34; *p*=6.4×10^−8^; variance explained mean model: 0.002%; variance explained variance model: 0.055%), whereas a region around rs10812921 on *LINGO2* was associated with ICV variability (chr9:28995582:C:A; MAF=0.4; *p*=5.5×10^−8^; variance explained variance model: 0.07%; variance explained mean model: 0.006%). Results were consistent when re-analyzing the data from healthy controls only (excluding participants with neuropsychiatric diagnoses): *p*=3.4×10^−11^ (rs59515793-pallidum), *p*=2.2×10^−8^ (rs6039642-cortical thickness), *p*=3.9×10^−8^ (rs9543733-thalamus) and *p*=1.4×10^−7^ (rs10812921-ICV). Figure 2 shows the relevant phenotype distributions for the top hits for the two models grouped by genotypes generated via the shift function ^16^. In short, the adopted shift function procedure was implemented in three stages: deciles of two phenotype distributions were calculated using the Harrell-Davis quantile estimator, followed by the computation of 95% confidence intervals of decile differences with bootstrap estimation of deciles’ standard error, and multiple comparison control so that the type I error rate remained close to 5% across the nine confidence intervals. Decile-by-decile shift function analysis confirmed reduce pallidum volume variance among homozygotes for the major rs59515793 allele (GG) in relation to the other two genotypes (GA, AA). Similarly, major allele homozygous subjects for rs6039642 (GG genotype) showed lower cortical thickness variance than carriers of the minor allele A. The rs10812921 heterozygotes and major allele homozygotes (AA, AC) had lower ICV variance than the participants with CC genotypes, whereas TC heterozygotes displayed higher thalamus variance than rs9543733 homozygotes (TT, CC).

**Figure 2.**
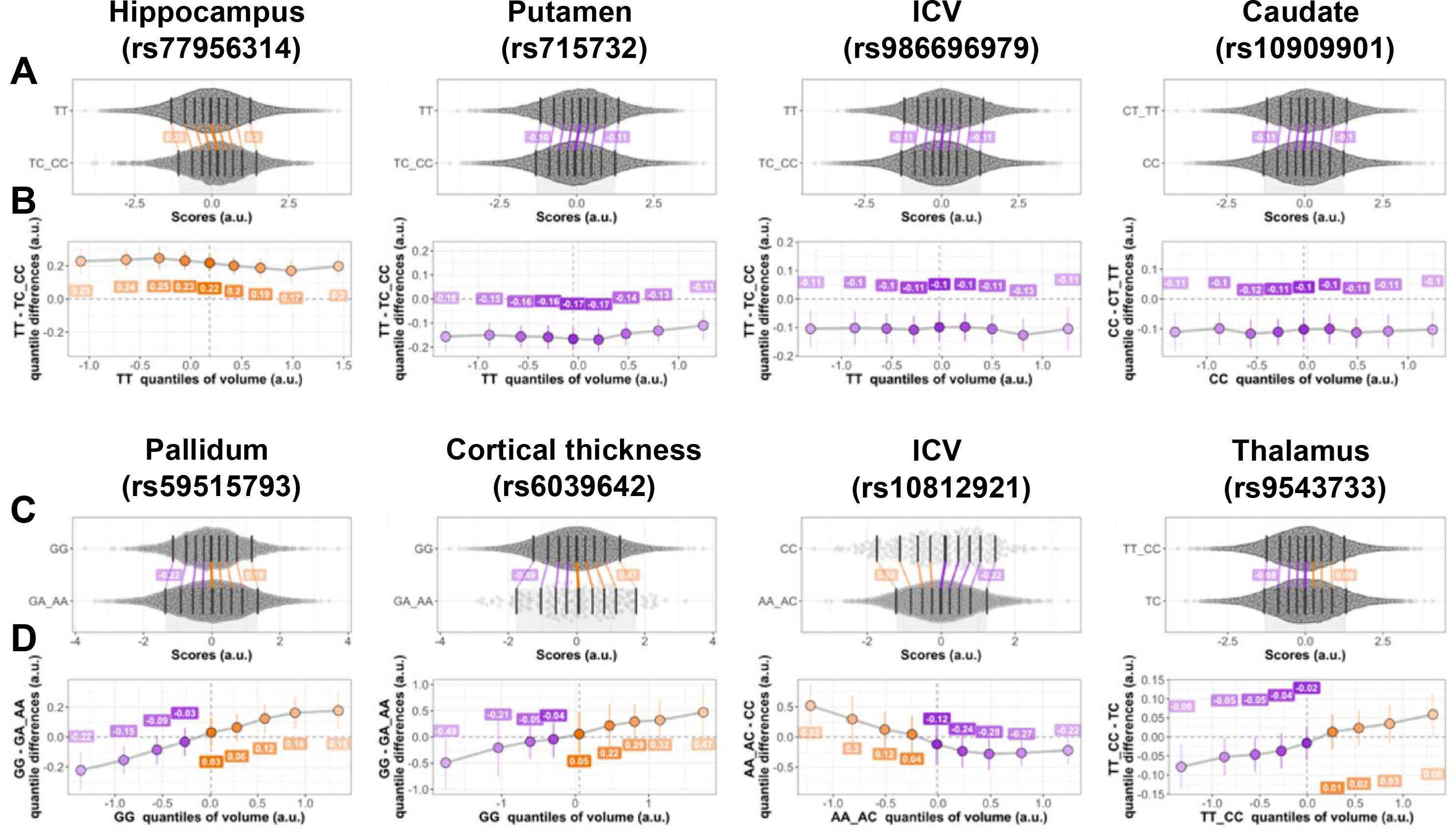
Shift function plots for the top genome-wide significant associations in mean and variance model GWAS results. The results corresponding to the top four mean models associations (conventional GWAS) are shown on the top rows (“A”, “B”), those corresponding to the top four variance model associations are displayed on the lower sections (“C”, “D”). A: Jittered marginal distribution scatterplots for the top four mean model associations, with overlaid shift function plots using deciles. Genotypes with the minor (effect) allele are shown as a single group. 95% confidence intervals were computed using a percentile bootstrap estimation of the standard error of the difference between quantiles on 1000 bootstrap samples. B: Linked deciles from shift functions on row “A”. C: Jittered marginal distribution scatterplots for the top four variance model associations, grouped by reference allele(s) versus effect allele(s) carriers. 95% confidence intervals were computed as in “A”. D: Linked deciles from shift functions on row “C”. For the mean model associations (“A” and “B”), variances explained by mean and variance parts of the model were 0.535% and 0.0006% (hippocampus, rs77956314), 0.444% and 0.0028% (putamen, rs715732), 0.211% and 0.0005% (ICV, rs986696979) and 0.254% and 0.0001% (caudate, rs10909901).

## Discussion

To our knowledge, this is the first evidence of genetic loci influencing variability of brain volumes beyond their mean value. A conceptually and methodologically similar approach revealed genetic control of the variance in body height and body mass index ^12^. Adding to the notion that phenotypic spread in a population is related to genetic variability, the current results show that the population variance of subcortical and intracranial volumes is partly under genetic control. Importantly, our findings on brain structure and the previous work on body mass index ^12^ provide converging evidence supporting the notion that common genetic variants affecting the mean and the variance of a trait need not be correlated and may influence phenotypes through complementary mechanisms.

Most variants associated with volumetric dispersion where at loci that have previously been linked to neuropsychiatric traits. Pallidum variability was related to a genotype near the neural cell adhesion molecule 2 gene (*NCAM2*), which has a documented role in neurodevelopment and has associations with Alzheimer’s disease and other neuropsychiatric phenotypes ^17–19^. Similarly, the significant variance locus for cortical thickness on chromosome 20 was located next to *PAK7* - a gene conferring risk for psychosis and involved in oxytocin gene networks of the brain ^20, 21^ - and the synaptosome associated protein 25 gene (*SNAP25*) - which participates in synaptic function and increases susceptibility for severe psychiatric conditions ^22, 23^. Moreover, the locus related to ICV variance was on *LINGO2*, which has been implicated in Parkinson disease and other psychiatric conditions ^24–26^.

Variance-controlling alleles can be interpreted as underlying distinct degrees of organismic robustness ^1^. Relevance to medical genetics also comes from the observation that several disease phenotypes emerge beyond a phenotypic threshold, which could be reached by the influence of high variability phenotypes ^27^. It is thus important to understand how the identified markers relate to brain variability under changing environments (robustness), how they interact with other genetic loci (epistasis) and how they relate to the clinical manifestation of disease. Similarly, variance-controlling loci can underlie variability from other genetic factors, potentially affecting evolutionary dynamics ^4^. Identifying the mechanisms by which variance-controlling genotypes influence gene expression variance in relevant brain structures may provide a proof of principle for the functional relevance of the identified genotypes. This type of effect on expression has been shown in model organisms ^28^, and the genomic loci identified here represent suitable candidates for targeted gene expression analysis in the human brain. The identification of specific genes involved in neural evolution and mental disorders suggests that brain variability in human populations is mediated by genetic factors. In so doing it also underscores the validity of gene-gene and gene-environment interactions in explaining heritability of complex human traits.

In summary, the results indicate that beyond associations with mean volumetric values, genotypic architecture modulates the variance of subcortical and intracranial dimensions across individuals. The lack of overlap between genetic associations detected by the standard additive genetic model and variance-controlling loci indicate independent mechanisms. These findings contribute to establish the genetic basis of phenotypic variance (i.e., heritability), allow identifying different degrees of brain robustness across individuals, and open new research avenues in the search for mechanisms controlling brain and mental health.

## Supplemental information

Supplemental information can be found with this article online.

## Author contributions

Conceptualization, ACP and LTW; Methodology, ACP and LTW; Investigation, ACP, vdM, TK; Writing – Original Draft, ACP and LTW; Writing – Review & Editing, all co-authors.

## Declaration of interest

The authors declare no competing interests.

## Data and code availability

All scripts are available upon reasonable request to the corresponding authors. Data availability notes for each cohort can be found on Supplementary Information.

## References

1. Félix MA, Barkoulas M. Pervasive robustness in biological systems. Nat Rev Genet 2015; 16(8):483–496.

2. Hibar DP, Stein JL, Renteria ME, Arias-Vasquez A, Desrivieres S, Jahanshad N et al. Common genetic variants influence human subcortical brain structures. Nature 2015; 520(7546):224–229.

3. Smeland OB, Wang Y, Frei O, Li W, Hibar DP, Franke B et al. Genetic Overlap Between Schizophrenia and Volumes of Hippocampus, Putamen, and Intracranial Volume Indicates Shared Molecular Genetic Mechanisms. Schizophr Bull 2018; 44(4):854–864.

4. Wagner A. Robustness and evolvability in living systems. Princeton University Press: Princeton, N.J.; Woodstock, 2005.

5. Masel J, Trotter MV. Robustness and evolvability. Trends Genet 2010; 26(9):406–414.

6. Starrfelt J, Kokko H. Bet-hedging––a triple trade-off between means, variances and correlations. Biol Rev Camb Philos Soc 2012; 87(3):742–755.

7. Shen X, Pettersson M, Rönnegård L, Carlborg Ö. Inheritance beyond plain heritability: variance-controlling genes in Arabidopsis thaliana. PLoS Genet 2012; 8(8):e1002839.

8. Chang CC, Chow CC, Tellier LC, Vattikuti S, Purcell SM, Lee JJ. Second-generation PLINK: rising to the challenge of larger and richer datasets. Gigascience 2015; 4:7.

9. R Core Team. R: A language and environment for statistical computing. R Foundation for Statistical Computing: Vienna, Austria, 2015.

10. Fischl B, Dale AM. Measuring the thickness of the human cerebral cortex from magnetic resonance images. Proceedings of the National Academy of Sciences of the United States of America 2000; 97(20):11050–11055.

11. Auer PL, Reiner AP, Leal SM. The effect of phenotypic outliers and non-normality on rare-variant association testing. Eur J Hum Genet 2016; 24(8):1188–1194.

12. Yang J, Loos RJ, Powell JE, Medland SE, Speliotes EK, Chasman DI et al. FTO genotype is associated with phenotypic variability of body mass index. Nature 2012; 490(7419):267–272.

13. Shen X, Carlborg O. Beware of risk for increased false positive rates in genome-wide association studies for phenotypic variability. Front Genet 2013; 4:93.

14. Watanabe K, Taskesen E, van Bochoven A, Posthuma D. Functional mapping and annotation of genetic associations with FUMA. Nat Commun 2017; 8(1):1826.

15. Franke B, Stein JL, Ripke S, Anttila V, Hibar DP, van Hulzen KJ et al. Genetic influences on schizophrenia and subcortical brain volumes: large-scale proof of concept. Nat Neurosci 2016; 19(3):420–431.

16. Rousselet GA, Pernet CR, Wilcox RR. Beyond differences in means: robust graphical methods to compare two groups in neuroscience. Eur J Neurosci 2017; 46(2):1738–1748.

17. Winther M, Berezin V, Walmod PS. NCAM2/OCAM/RNCAM: cell adhesion molecule with a role in neuronal compartmentalization. Int J Biochem Cell Biol 2012; 44(3):441–446.

18. Petit F, Plessis G, Decamp M, Cuisset JM, Blyth M, Pendlebury M et al. 21q21 deletion involving NCAM2: report of 3 cases with neurodevelopmental disorders. Eur J Med Genet 2015; 58(1):44–46.

19. Leshchyns’ka I, Liew HT, Shepherd C, Halliday GM, Stevens CH, Ke YD et al. Aβ-dependent reduction of NCAM2-mediated synaptic adhesion contributes to synapse loss in Alzheimer’s disease. Nat Commun 2015; 6:8836.

20. Morris DW, Pearson RD, Cormican P, Kenny EM, O’Dushlaine CT, Perreault LP et al. An inherited duplication at the gene p21 Protein-Activated Kinase 7 (PAK7) is a risk factor for psychosis. Hum Mol Genet 2014; 23(12):3316–3326.

21. Quintana DS, Rokicki J, van der Meer D, Alnæs D, Kaufmann T, Córdova-Palomera A et al. Oxytocin pathway gene networks in the human brain. Nat Commun 2019; 10(1):668.

22. Houenou J, Boisgontier J, Henrion A, d’Albis MA, Dumaine A, Linke J et al. A Multilevel Functional Study of a. J Neurosci 2017; 37(43):10389–10397.

23. Najera K, Fagan BM, Thompson PM. SNAP-25 in Major Psychiatric Disorders: A Review. Neuroscience 2019.

24. Williams SM, An JY, Edson J, Watts M, Murigneux V, Whitehouse AJO et al. An integrative analysis of non-coding regulatory DNA variations associated with autism spectrum disorder. Mol Psychiatry 2018.

25. Wu YW, Prakash KM, Rong TY, Li HH, Xiao Q, Tan LC et al. Lingo2 variants associated with essential tremor and Parkinson’s disease. Hum Genet 2011; 129(6):611–615.

26. Vilariño-Güell C, Wider C, Ross OA, Jasinska-Myga B, Kachergus J, Cobb SA et al. LINGO1 and LINGO2 variants are associated with essential tremor and Parkinson disease. Neurogenetics 2010; 11(4):401–408.

27. Ayroles JF, Buchanan SM, O’Leary C, Skutt-Kakaria K, Grenier JK, Clark AG et al. Behavioral idiosyncrasy reveals genetic control of phenotypic variability. Proc Natl Acad Sci U S A 2015; 112(21):6706–6711.

28. Huang W, Carbone MA, Magwire MM, Peiffer JA, Lyman RF, Stone EA et al. Genetic basis of transcriptome diversity in Drosophila melanogaster. Proc Natl Acad Sci U S A 2015; 112(44):E6010–6019.

