## Supplementary Information for "Genetic control of variability in subcortical and intracranial volumes"

**Title:** Genetic control of phenotypic variability in subcortical and intracranial volumes

| **Cohort** | **Source** | **Comment** | **References** |
| --- | --- | --- | --- |
| BETULA | http://www.org.umu.se/betula/betula/?languageId=1 | Betula was supported by a Wallenberg Scholar Grant (KAW). | (Nilsson et al., 2004) |
| HUBIN | Authors | This study was supported by the Swedish Research Council (2006-2992, 2006-986, K2007-62X-15077-04-1, 2008-2167, K2008-62P-20597-01-3. K2010-62X-15078-07-2, K2012-61X-15078-09-3, 2017-00949), the regional agreement on medical training and clinical research between Stockholm County Council and the Karolinska Institutet, the Knut and Alice Wallenberg Foundation, and the HUBIN project. | (Haukvik et al., 2012) |
| HUNT | https://www.ntnu.edu/hunt | The HUNT Study is a collaboration between HUNT Research Centre, Faculty of Medicine and Health Sciences, Norwegian University of Science and Technology (NTNU), Nord-Trøndelag County Council, Central Norway Regional Health Authority, and the Norwegian Institute of Public Health. HUNT-MRI and the genetic analysis were funded by grants from the Liaison Committee between the Central Norway Regional Health Authority and NTNU to principal investigator Asta Håberg, and the Norwegian National Advisory Unit for functional MRI. We thank the HUNT MRI participants, MRI technicians and the Department of Diagnostic Imaging at Levanger Hospital, Professor Lars Jacob Stovner (NTNU) and the administrative staff at HUNT. | (Honningsvåg et al., 2012; Håberg et al., 2016) |
| NCNG | Authors | The sample collection was supported by grants from the Bergen Research Foundation and the University of Bergen, the Dr Einar Martens Fund, the K.G. Jebsen Foundation, the Research Council of Norway, to SLH, VMS, and TE. | (Espeseth et al., 2012) |
| NIMAGE | [www.neuroimage.nl](http://www.neuroimage.nl) | This project was supported by grants from National Institutes of Health (grant R01MH62873 to SV Faraone) for initial sample recruitment, and from NWO Large Investment (grant 1750102007010 to JK Buitelaar), NWO Brain & Cognition (grant 433-09-242 to JK Buitelaar), ZonMW Grant 60-60600-97-193, and grants from Radboud University Medical Center, University Medical Center Groningen, Accare, and VU University Amsterdam for subsequent assessment waves. NeuroIMAGE also receives funding from the European Community’s Seventh Framework Programme (FP7/2007 – 2013) under grant agreements n° 602805 (Aggressotype), n° 278948 (TACTICS), and n° 602450 (IMAGEMEND), and from the European Community’s Horizon 2020 Programme (H2020/2014 – 2020) under grant agreements n° 643051 (MiND) and n° 667302 (CoCA). | (von Rhein et al., 2015) |
| PNC | https://www.med.upenn.edu | Support for the collection of the data sets was provided by the National Institutes of Health (grant RC2MH089983 awarded to R. Gur and grant RC2MH089924 awarded to H. Hakonarson). Subjects were recruited through the Center for Applied Genomics at The Children’s Hospital in Philadelphia. | (Satterthwaite et al., 2016; Satterthwaite et al., 2014) |
| STROKEMRI | Authors | Supported by the Research Council of Norway (249795, 248238), the South-Eastern Norway Regional Health Authority (2014097, 2015044, 2015073, 2016083), and the Norwegian ExtraFoundation for Health and Rehabilitation (2015/FO5146). | (Dørum et al., 2016) |
| TOP | Authors | The work was funded by the Research Council of Norway (213837, 223273, 204966/F20, 213694, 229129, 249795/F20, 248778), the South-Eastern Norway Regional Health Authority (2013-123, 2014-097, 2015-073, #2017-112) and Stiftelsen Kristian Gerhard Jebsen. | (Brandt et al., 2015; Kaufmann et al., 2017; Kaufmann et al., 2015; Skåtun et al., 2016) |
| UBA | Authors | Swiss National Science Foundation (grants 163434, 147570 and 159740) | (Heck et al., 2014) |
| UKBB | https://www.ukbiobank.ac.uk/ | All subjects with a primary or secondary ICD-10 diagnosis with a mental or neurological disorder were excluded prior to analysis and the remaining subjects included as healthy controls. The used UK Biobank project ID number is #27412. | (Alfaro-Almagro et al., 2018) |
| UNIBA | Authors | This work was supported by a “Capitale Umano ad Alta Qualificazione” grant by Fondazione Con Il Sud awarded to Alessandro Bertolino and by a Hoffmann-La Roche Collaboration Grant awarded to Giulio Pergola. This paper reflects only the author's views and the European Union is not liable for any use that may be made of the information contained therein. | (Pergola et al., 2017) |

**Supplementary Table S1.** Additional details on data collection, and sample-specific acknowledgements

**Supplementary Methods**

*Quality control and pre-processing of brain scans*

A harmonized analysis protocol was applied to all individual subject raw data, which were stored and analysed locally at the University of Oslo. Surface-based morphometry and subcortical segmentation were performed using automated pipelines from FreeSurfer v5.3. Following standard protocols (<https://surfer.nmr.mgh.harvard.edu/fswiki/Edits>), several samples were screened by trained research personnel to detect segmentation errors, evaluate the quality of each brain scan and make segmentation edits where possible. When appropriate, bad quality images were discarded. Nevertheless, visual inspection and manual editing of each image was not possible due to the large number of individuals in the study; therefore, an automated quality control pipeline was applied to exclude potential outliers based on global data quality measures (in addition to the described manual edits/exclusions). Briefly, age, age², sex and scanning site were regressed out from mean cortical thickness, cortex volume, subcortical grey matter volume and from estimated total intracranial volume; then, the absolute of the residuals was z-standardized and subjects that exceeded a pre-defined 4-standard deviation threshold were excluded. Overall, images were excluded either based on manual QC or because they exceeded the standard deviation limit.

*Quality control and imputation of genetic data*

Genotyping was conducted at each site using commercially available genotyping platforms. For all cohorts except UK Biobank (UKBB), phasing and imputation were performed in-house following protocols in line with those applied by the ENIGMA consortium (http://enigma.ini.usc.edu). Briefly, before imputation, markers were excluded based on genotyping missingness (above 5%), minor allele frequency (MAF) under 1% or deviating from Hardy Weinberg equilibrium (*p*<10^-6^). Individuals with high rates of genotyping missingness (above 5%), cryptic relatedness (pi-hat above 18.5%) or genome-wide heterozygosity (beyond mean ±4 SD of the sample) were removed from the analyses. Analyses were restricted to individuals from European ancestry as determined through multidimensional scaling. Then, imputation was performed using MACH (<http://www.sph.umich.edu/csg/abecasis/MACH>) onto reference haplotypes from the 1000 Genomes Project (build 37, assembly hg19). After imputation, genotypes were additionally screened to remove SNPs with low imputation quality (estimated pseudo-*R*^2^<0.3), low MAF (<5%) or failing HWE (*p*<10^-6^). For the UKBB, we used the provided imputed data, processed with established protocols, and carried out the same post-imputation quality control steps as described for the other samples.

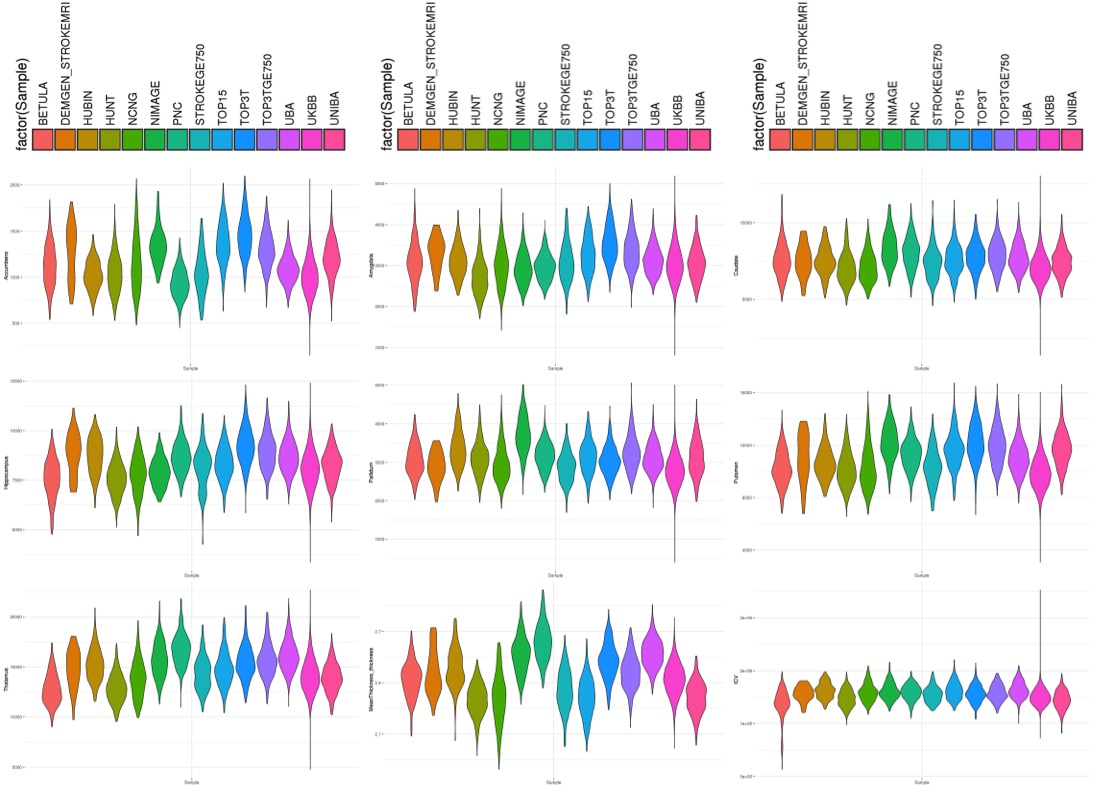

**Supplementary Figure S1.** Distribution of volumetric measurements across samples, for the nine included traits

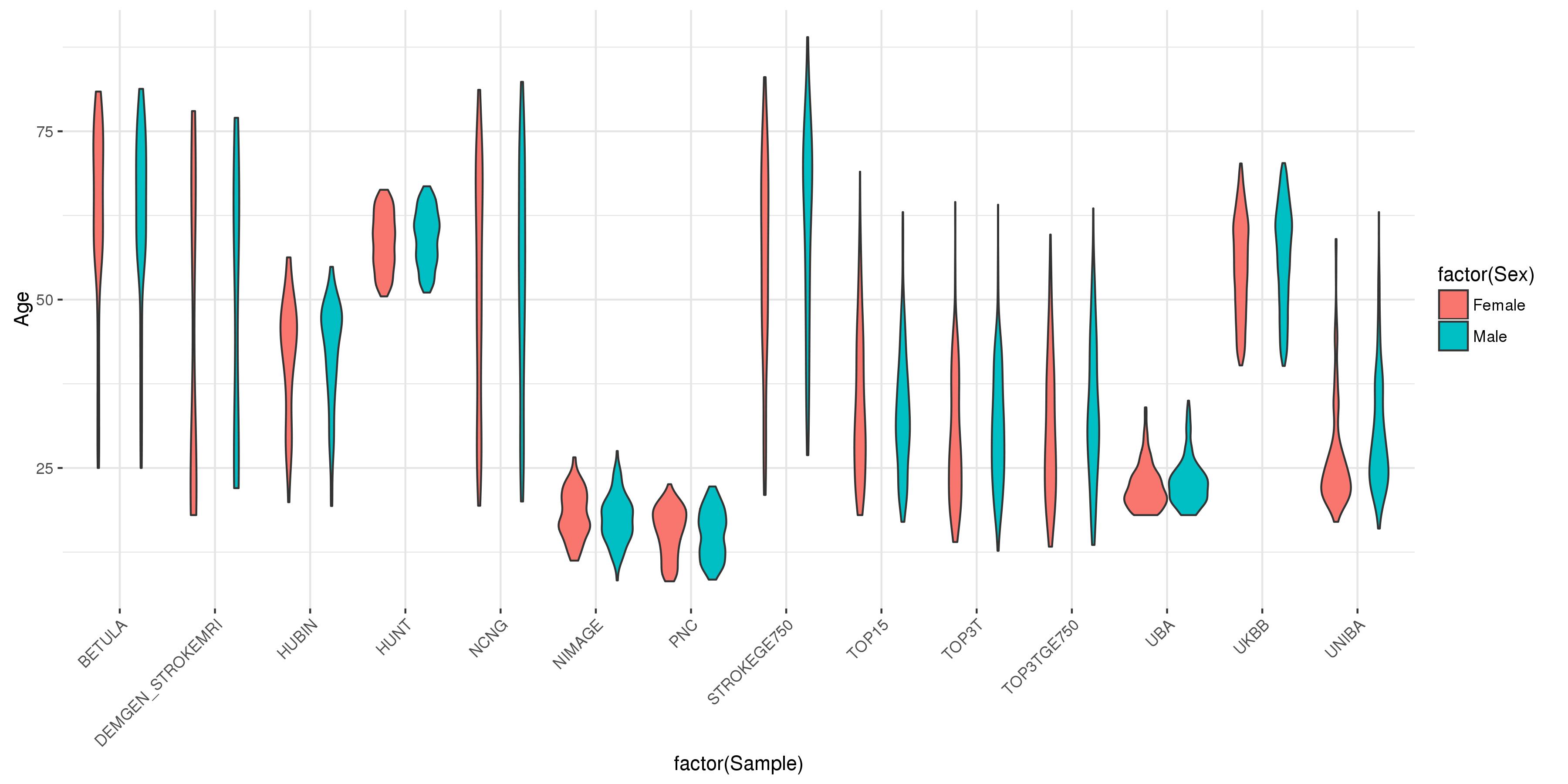

**Supplementary Figure S2.** Distribution of sex and age demographics across samples

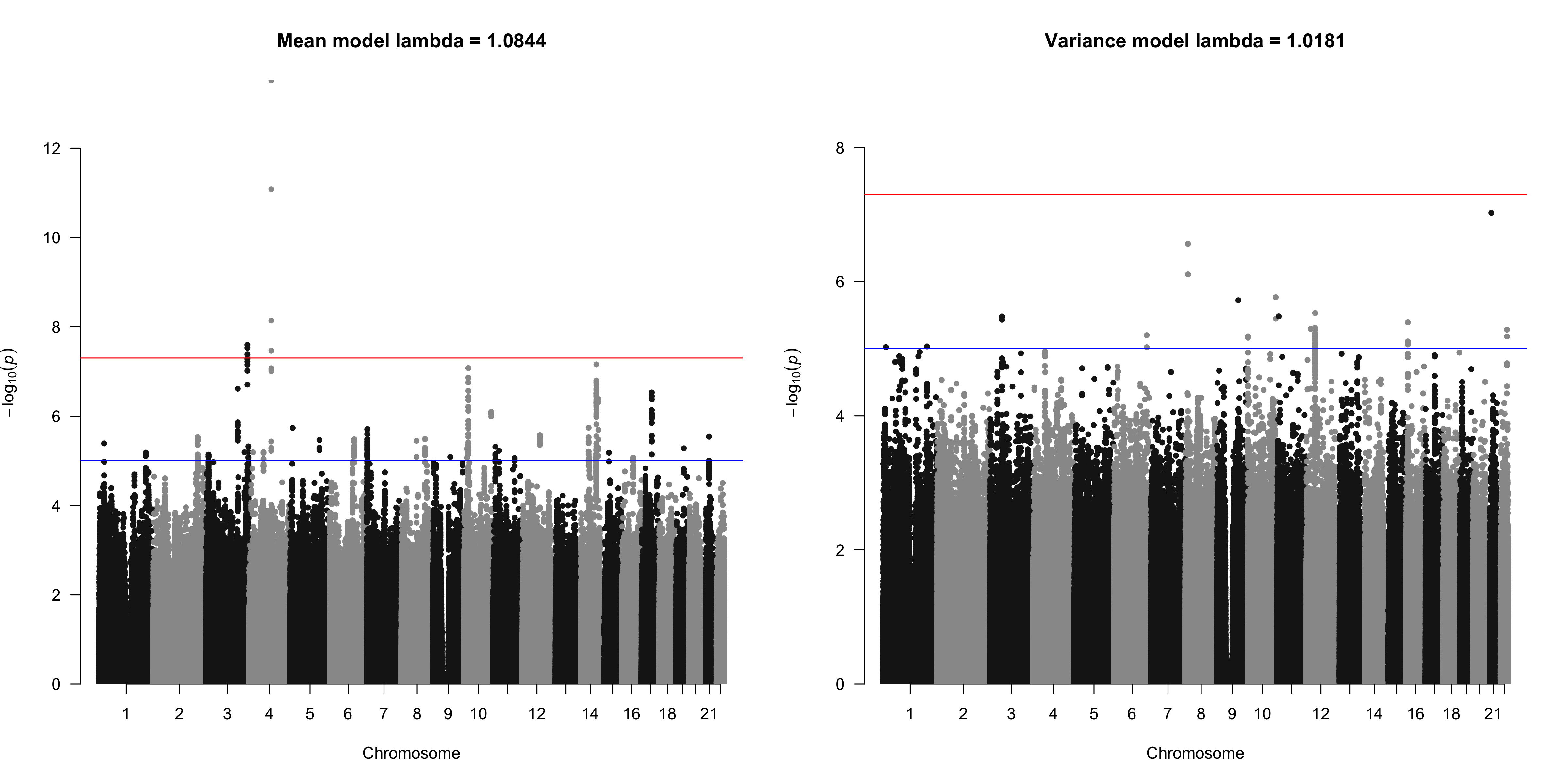

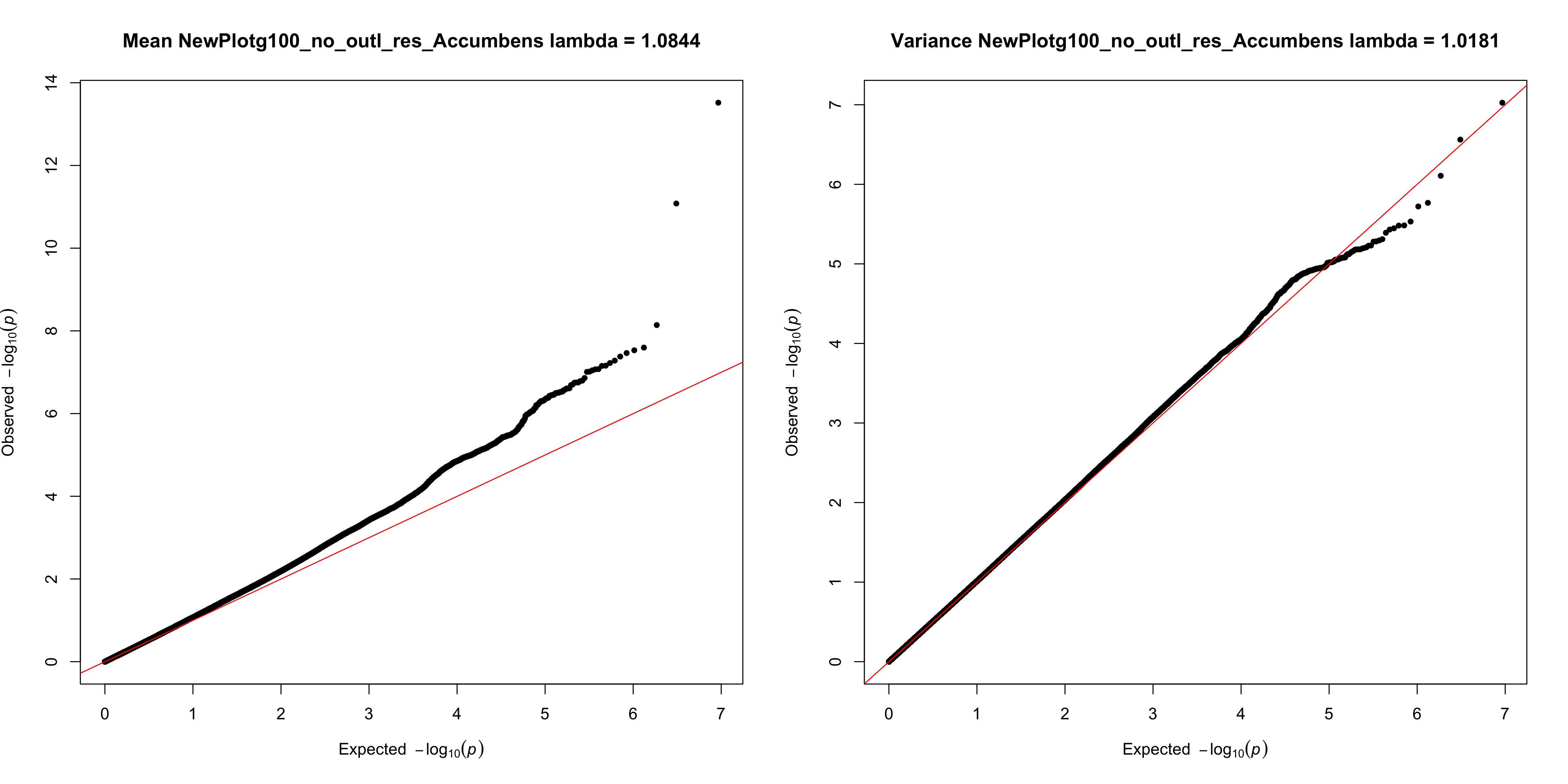

**Supplementary Figure S3a.** Manhattan and QQ plots for unadjusted mean- and variance-GWAS results, before genomic inflation correction (Accumbens)

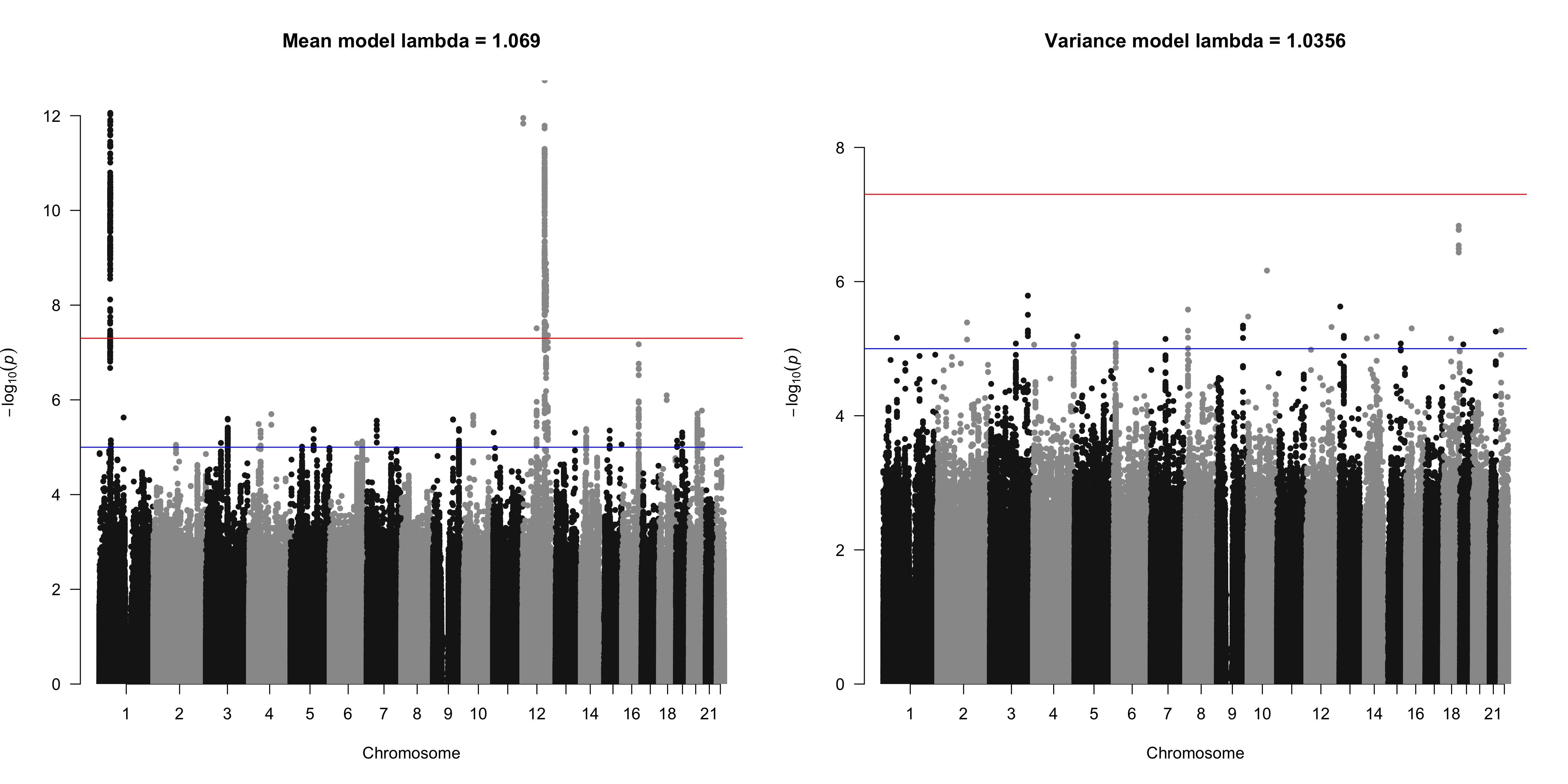

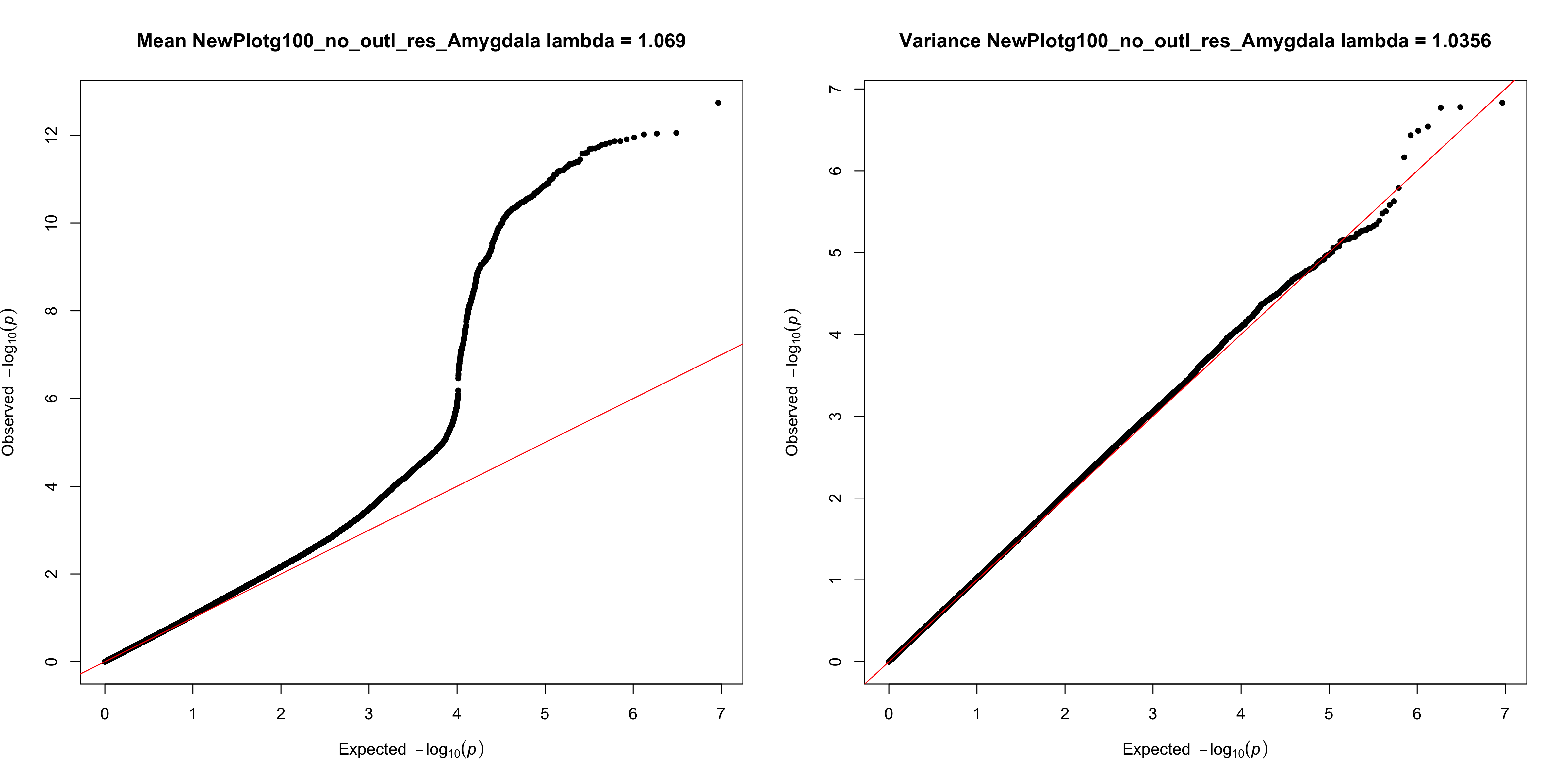
**Supplementary Figure S3b.** Manhattan and QQ plots for unadjusted mean- and variance-GWAS results, before genomic inflation correction (Amygdala)

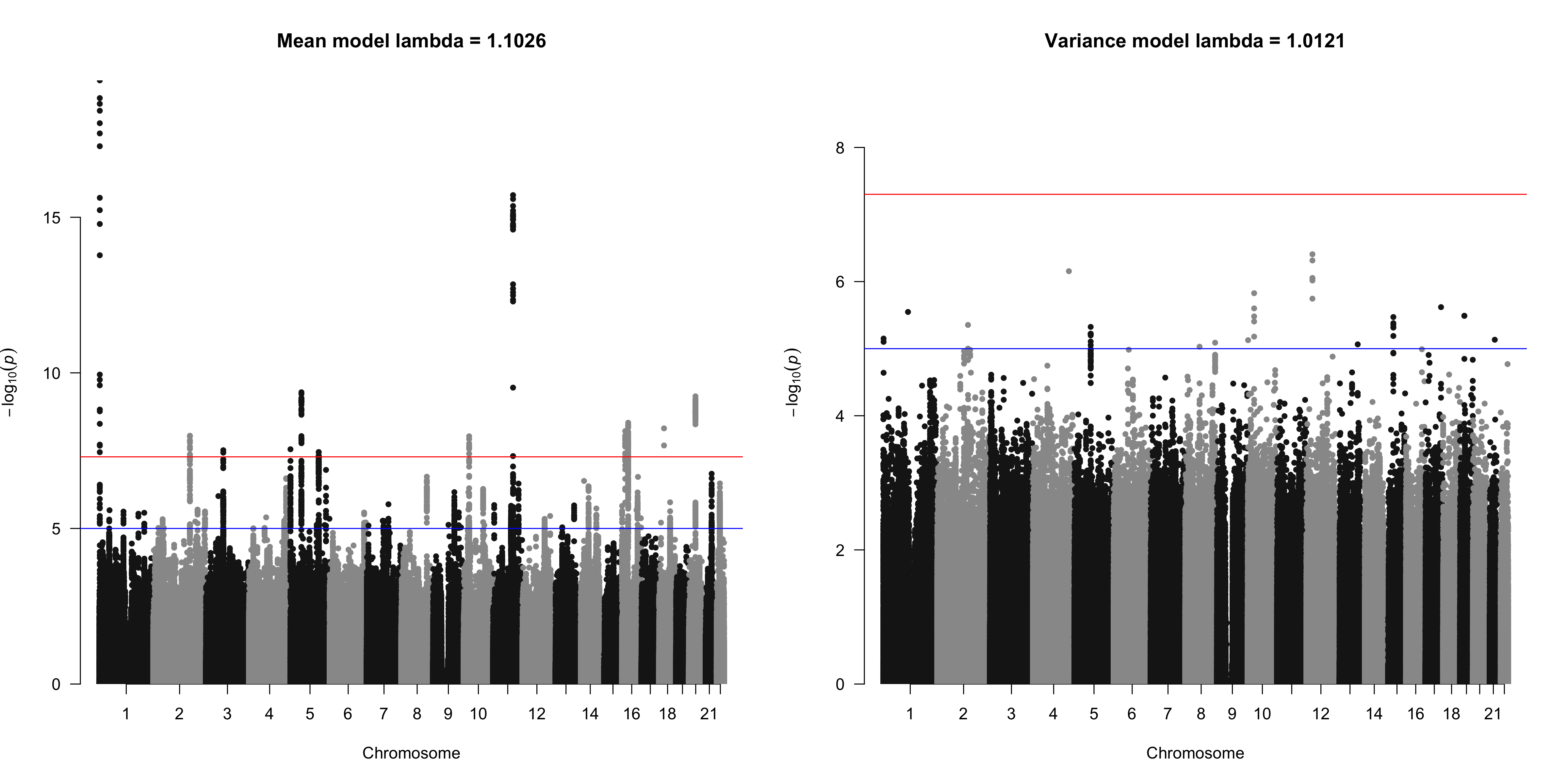

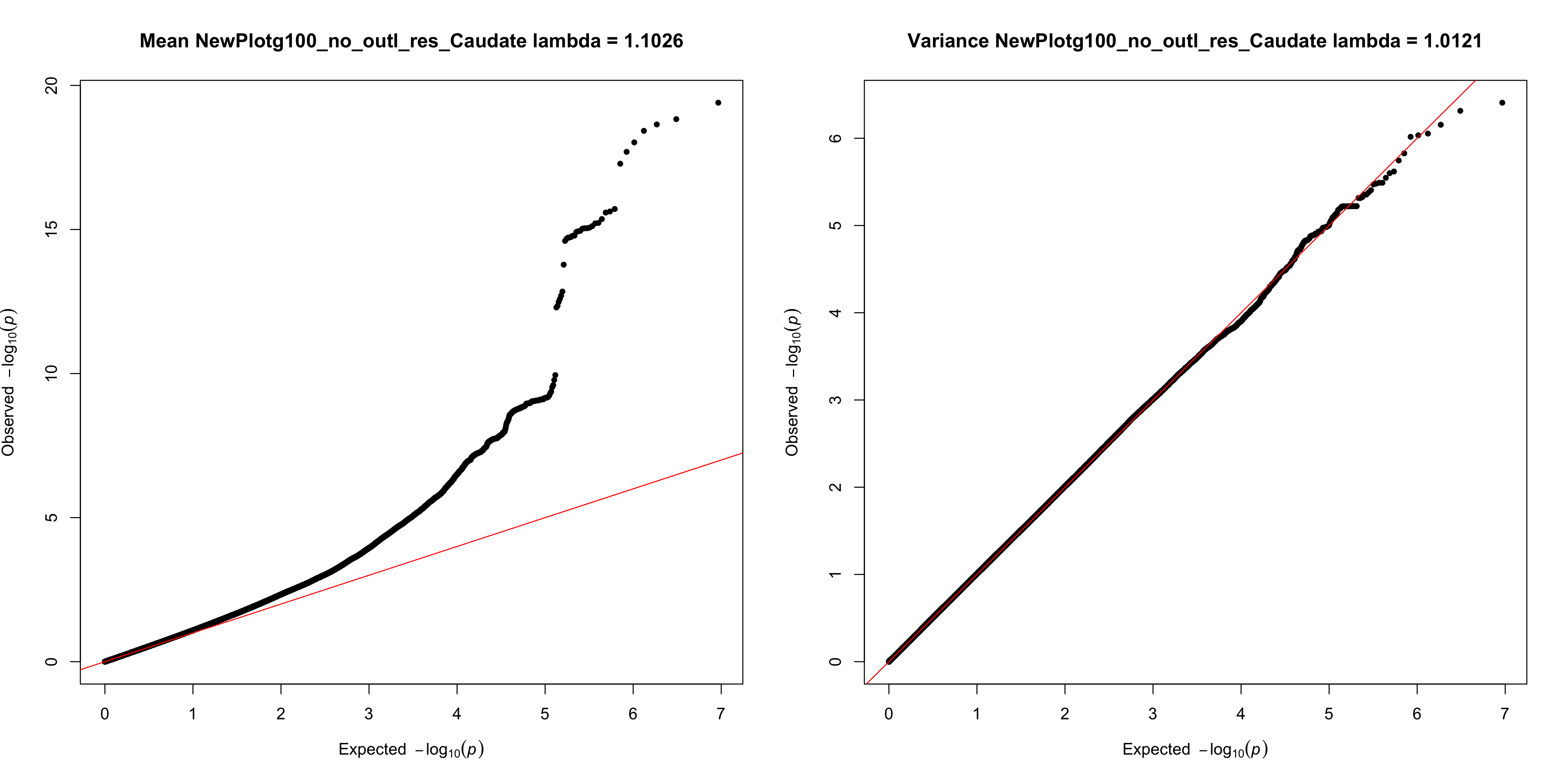
**Supplementary Figure S3c.** Manhattan and QQ plots for unadjusted mean- and variance-GWAS results, before genomic inflation correction (Caudate)

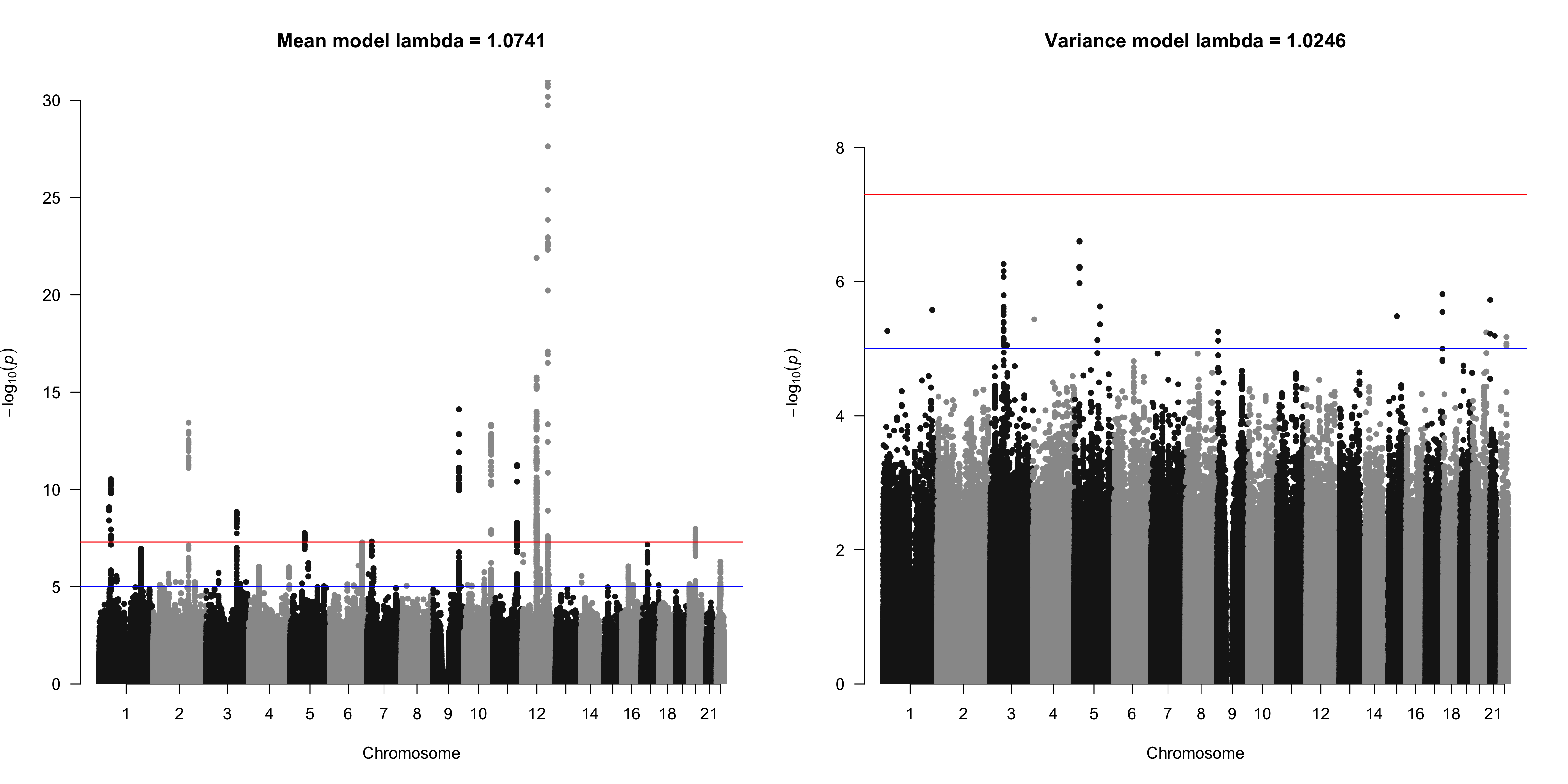

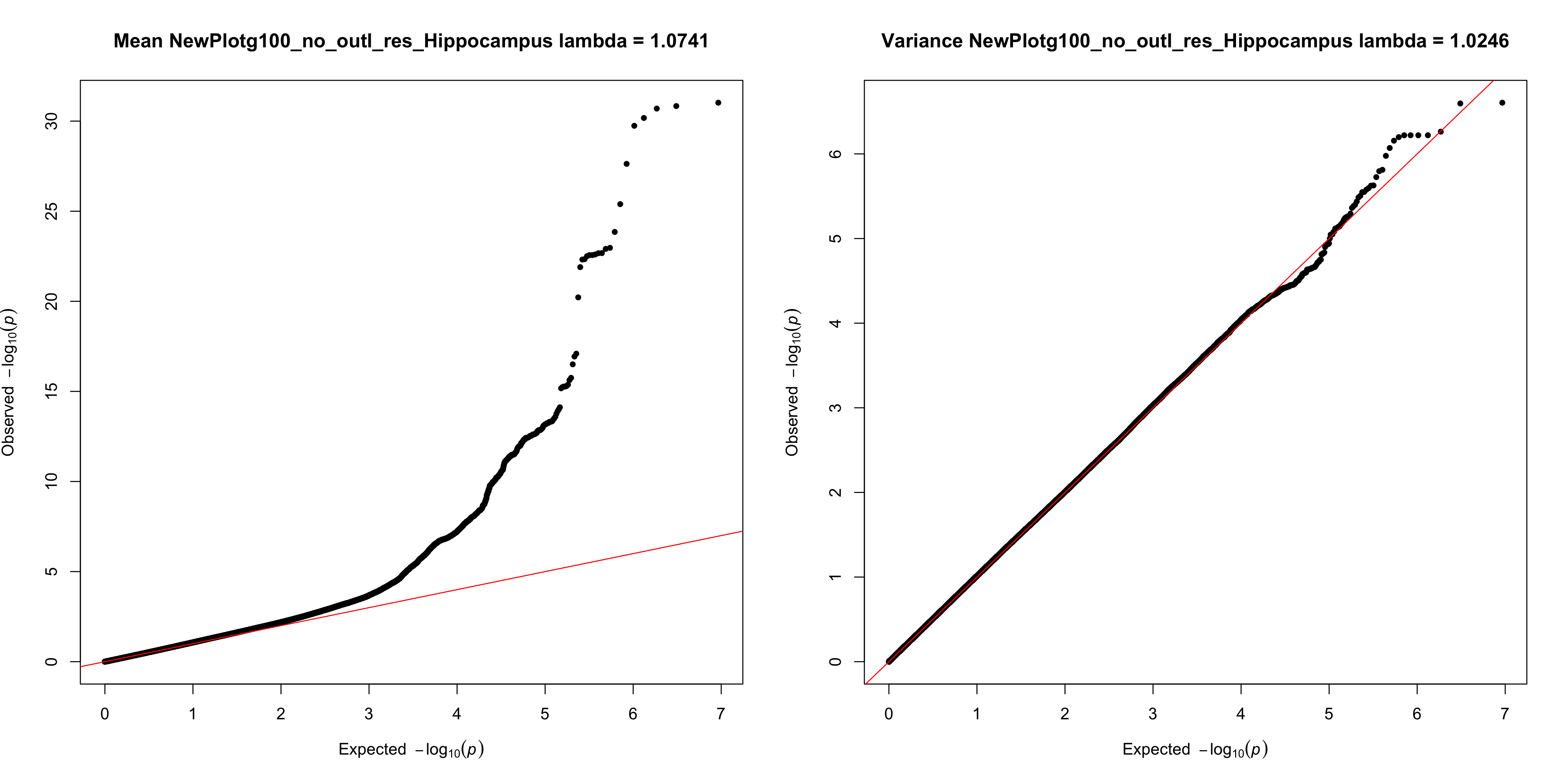
**Supplementary Figure S3d.** Manhattan and QQ plots for unadjusted mean- and variance-GWAS results, before genomic inflation correction (Hippocampus)

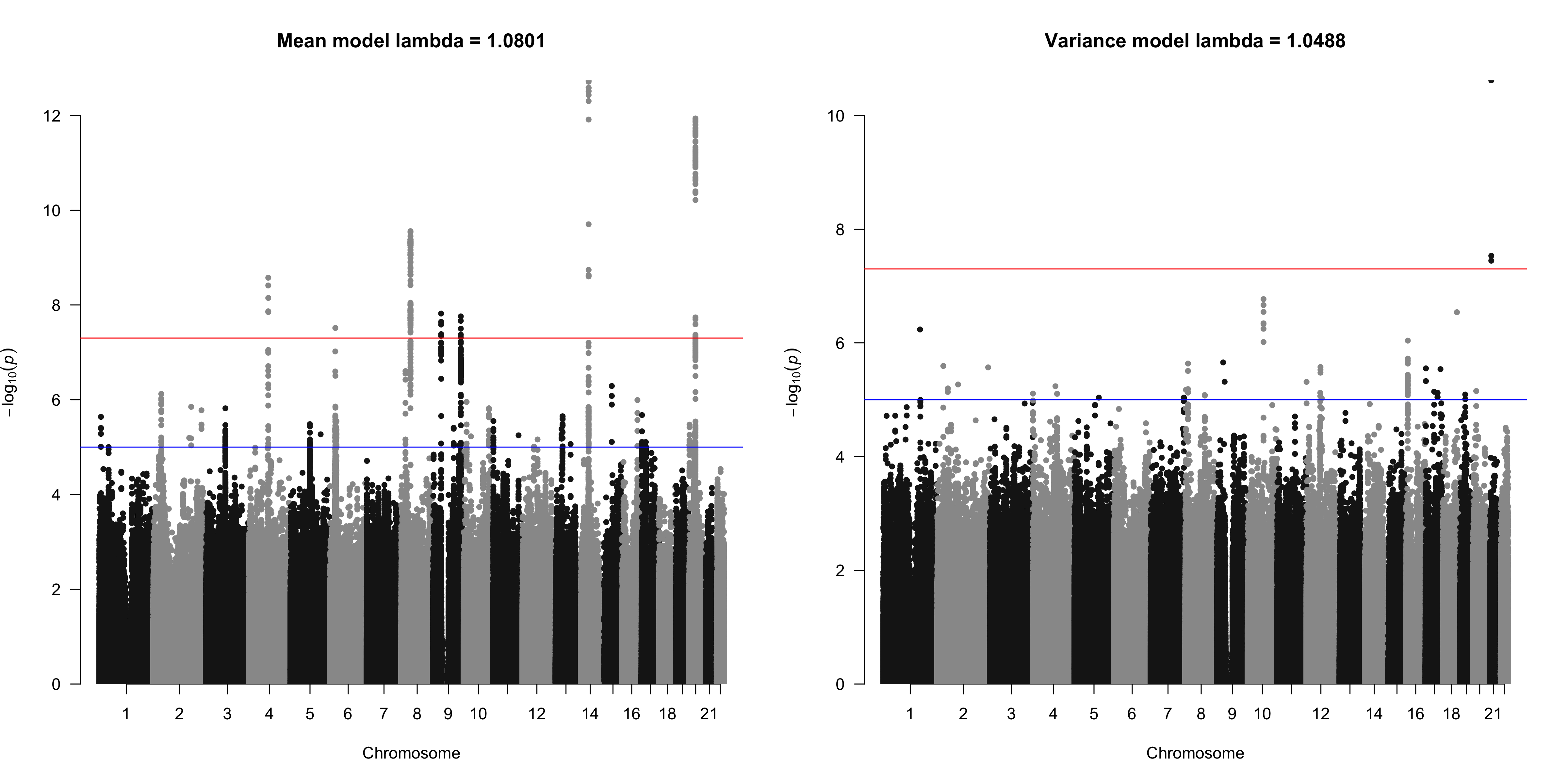

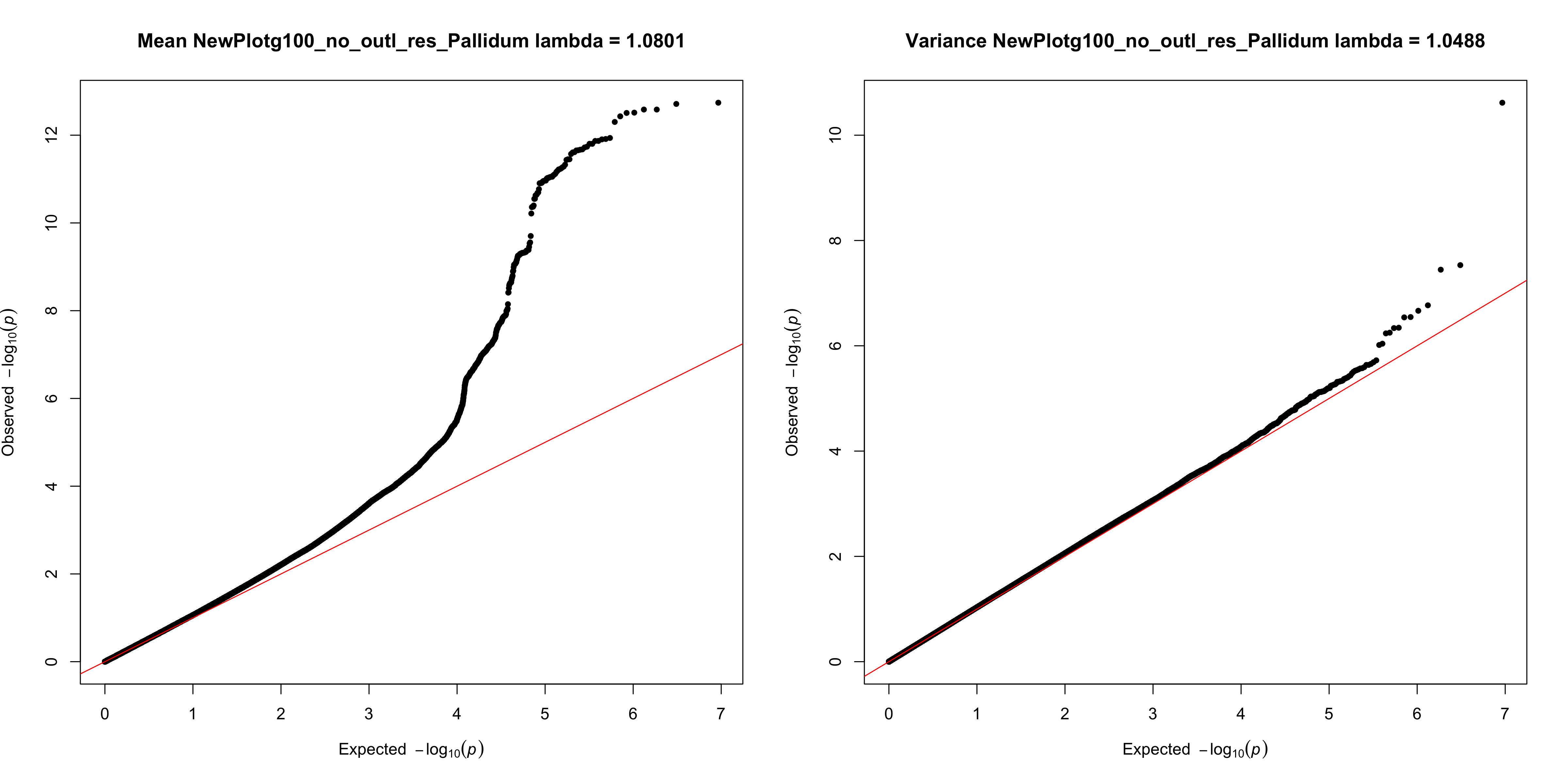
**Supplementary Figure S3e.** Manhattan and QQ plots for unadjusted mean- and variance-GWAS results, before genomic inflation correction (Pallidum)

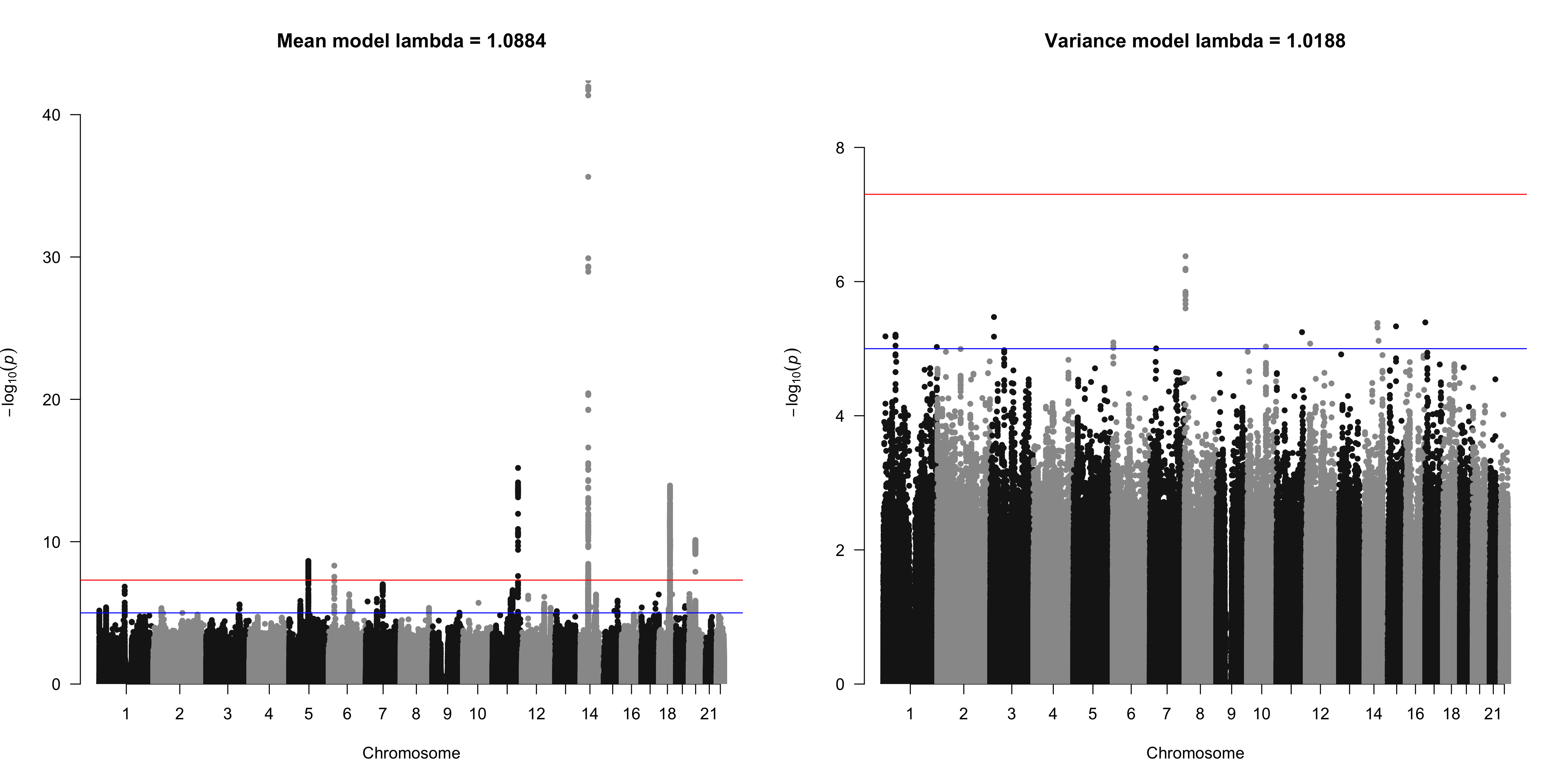

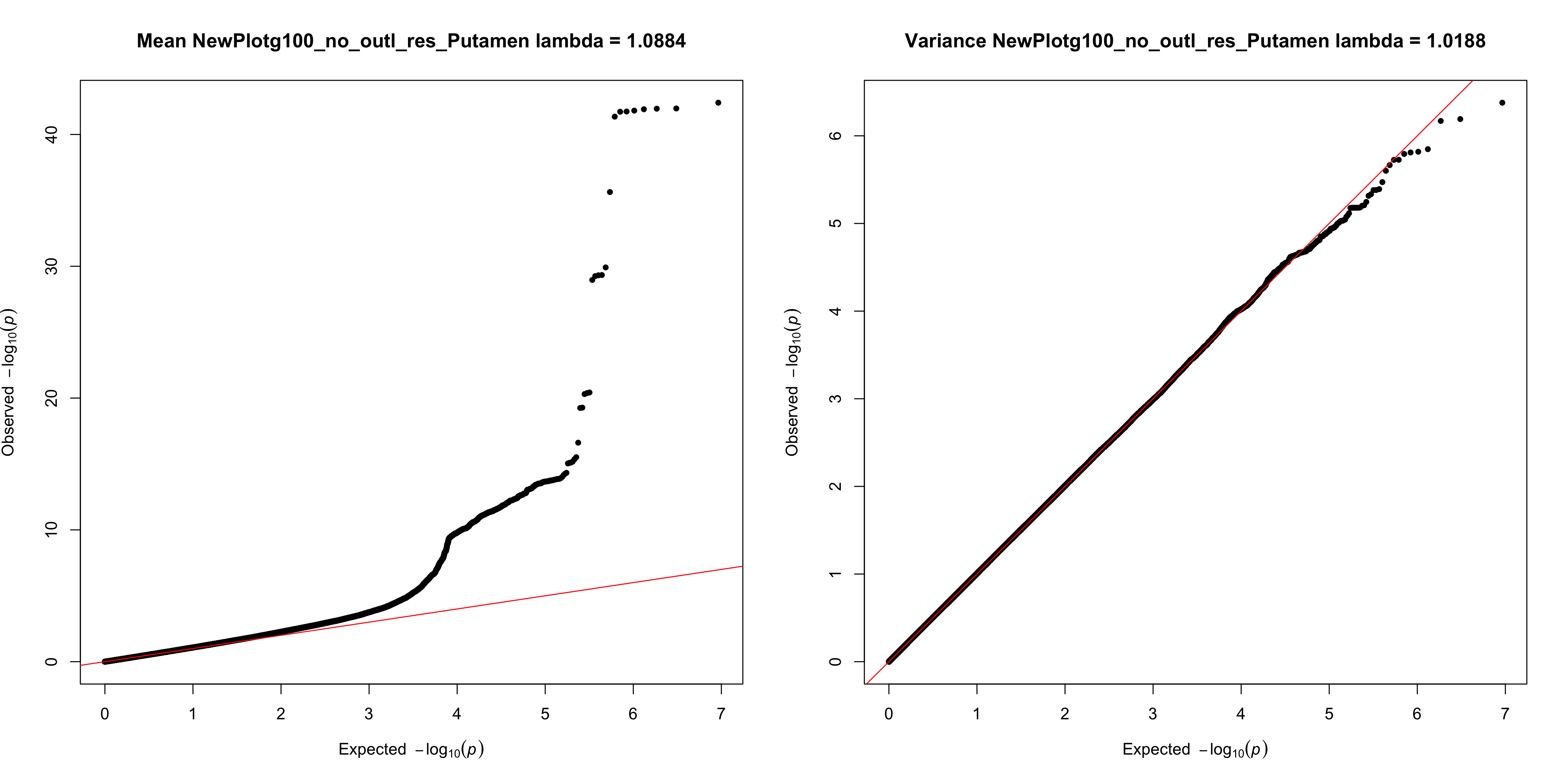
**Supplementary Figure S3f.** Manhattan and QQ plots for unadjusted mean- and variance-GWAS results, before genomic inflation correction (Putamen)

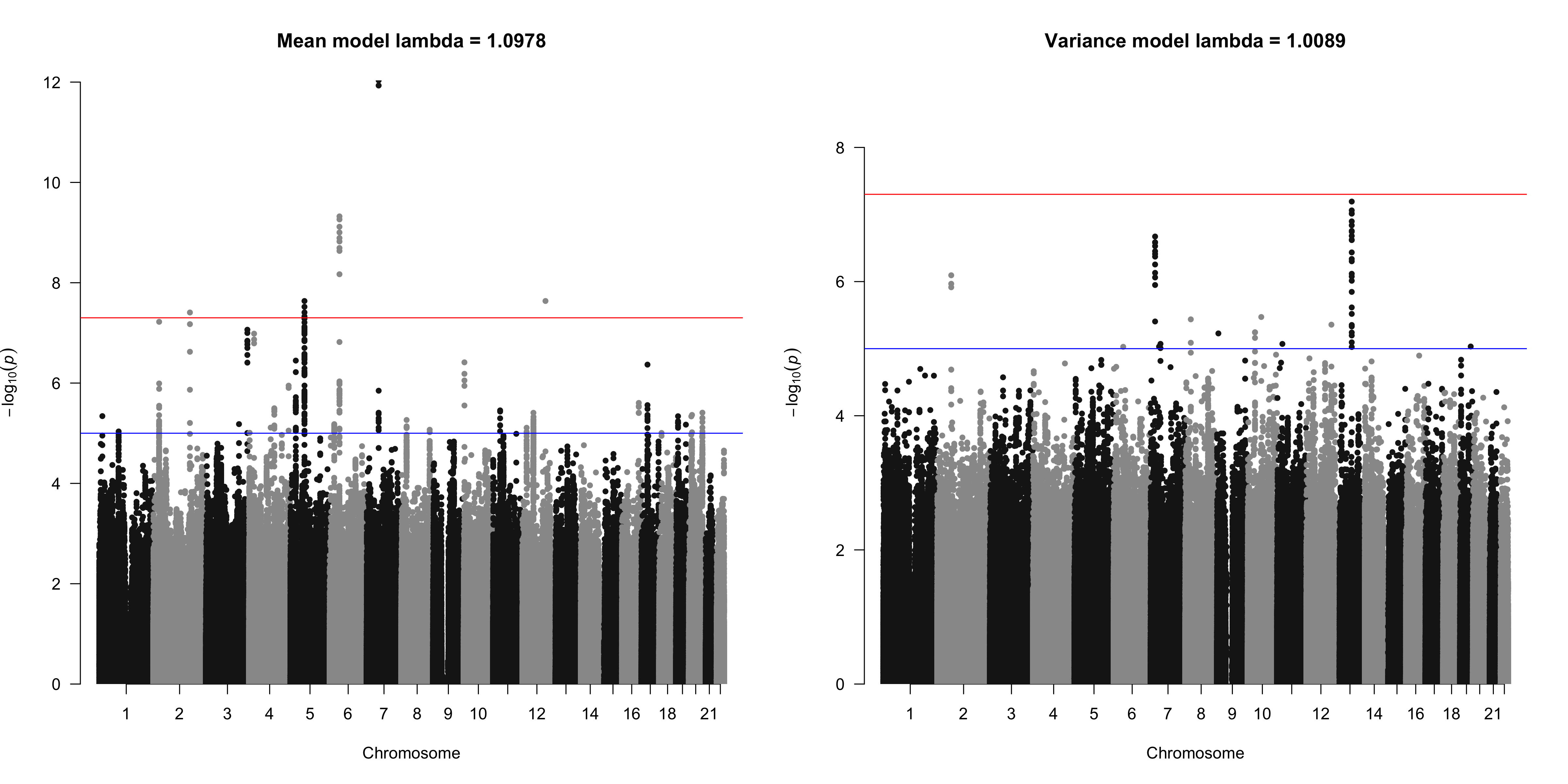

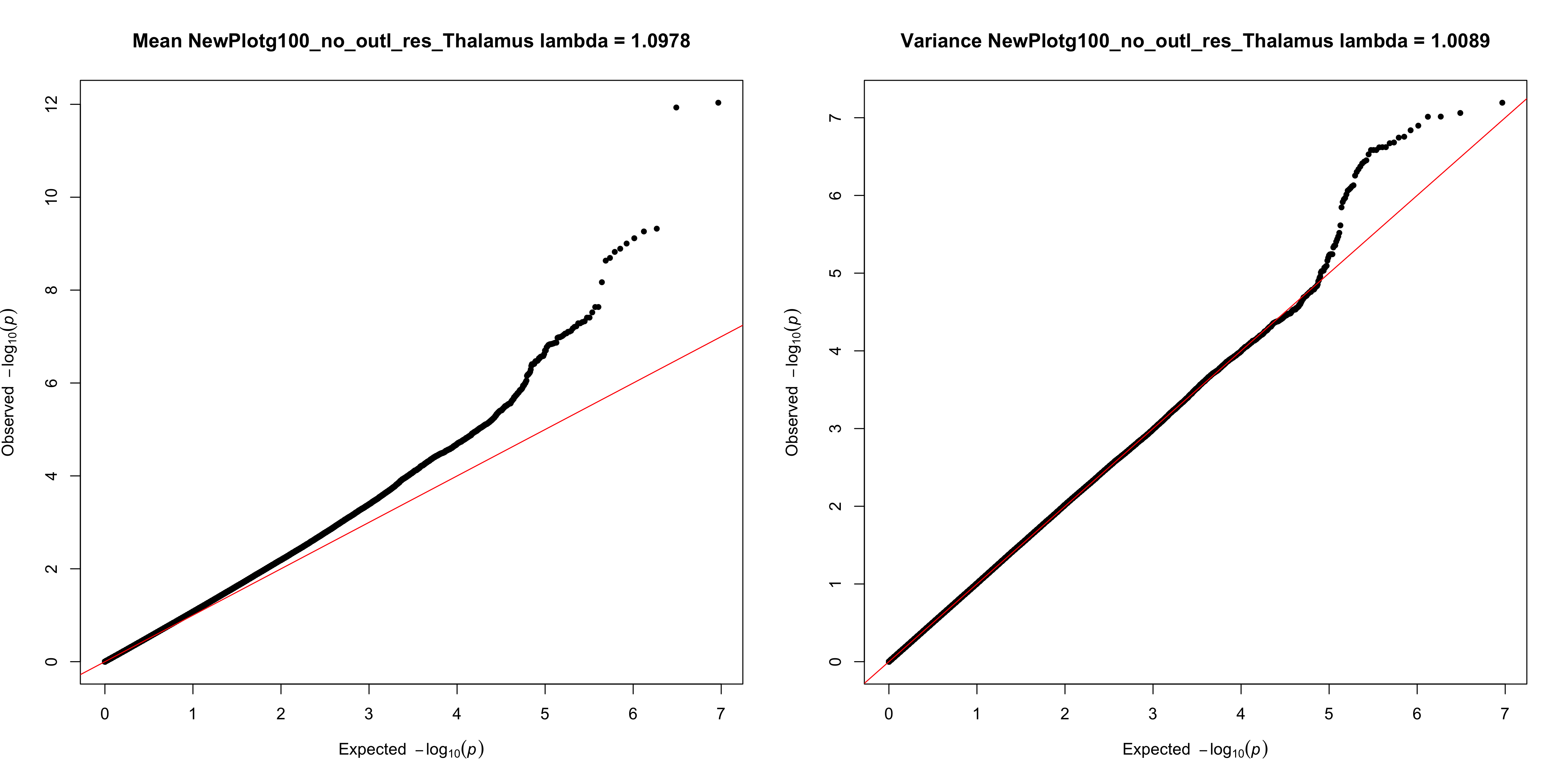
**Supplementary Figure S3g.** Manhattan and QQ plots for unadjusted mean- and variance-GWAS results, before genomic inflation correction (Thalamus)

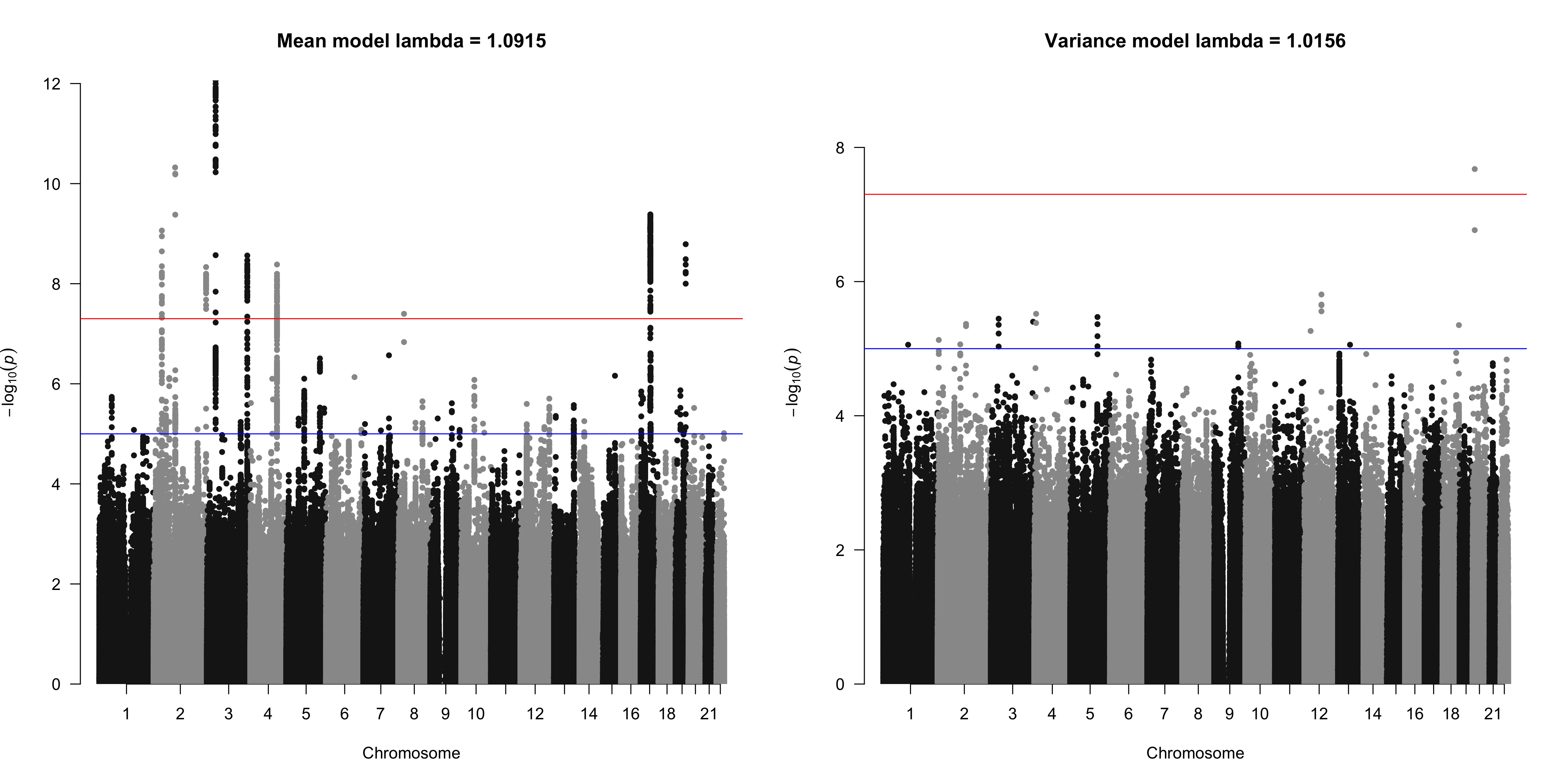

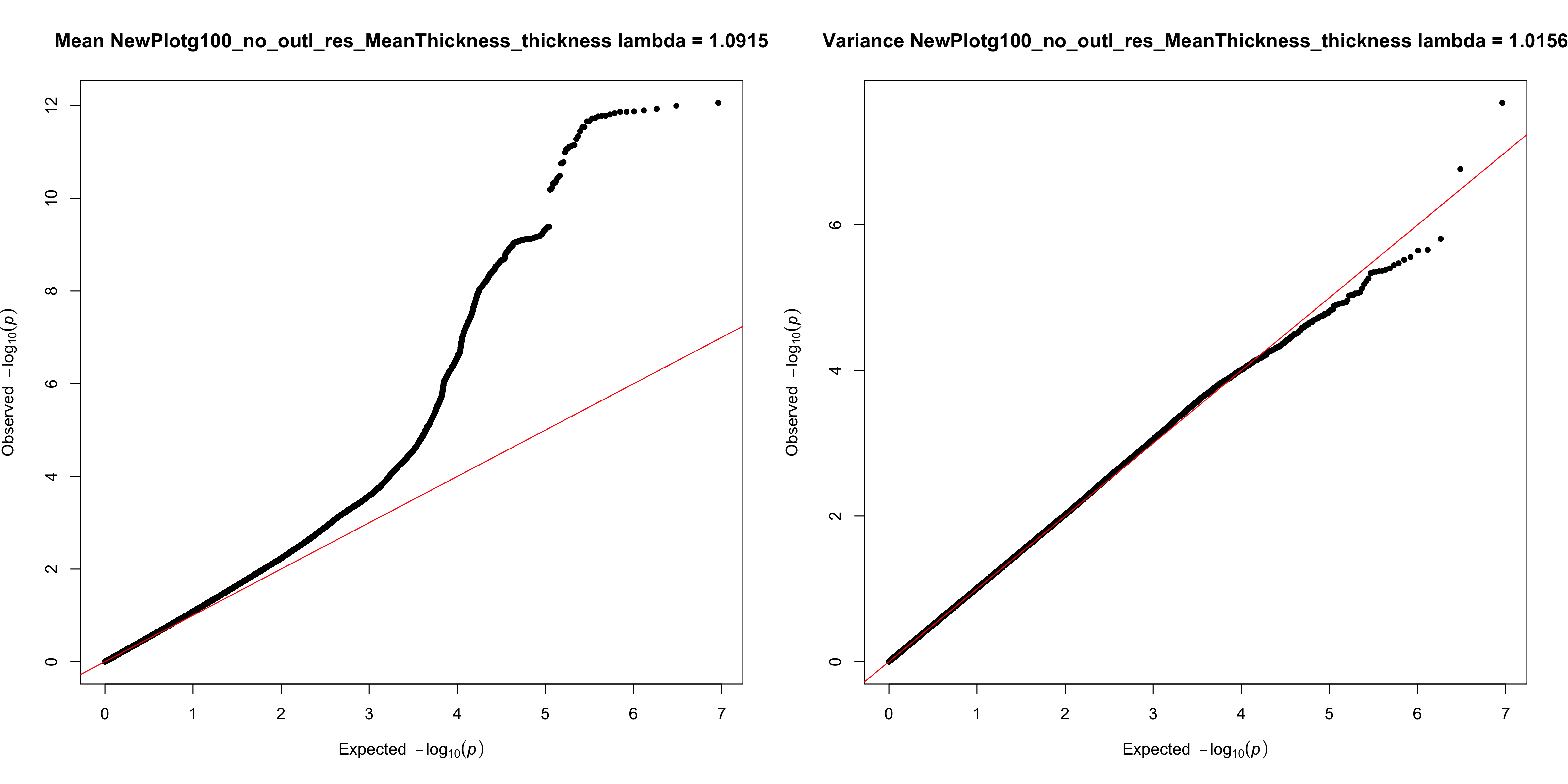
**Supplementary Figure S3h.** Manhattan and QQ plots for unadjusted mean- and variance-GWAS results, before genomic inflation correction (mean cortical thickness)

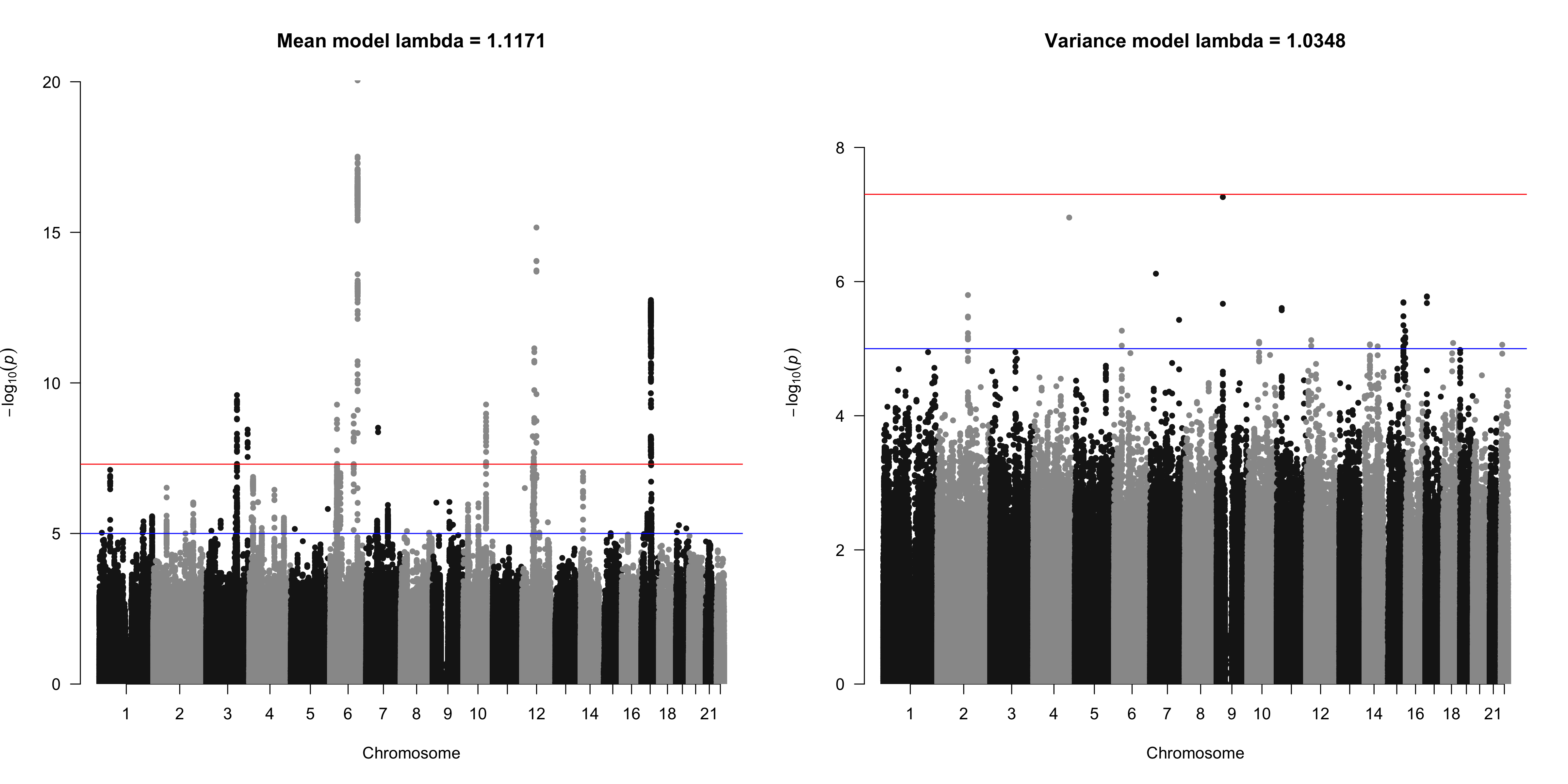

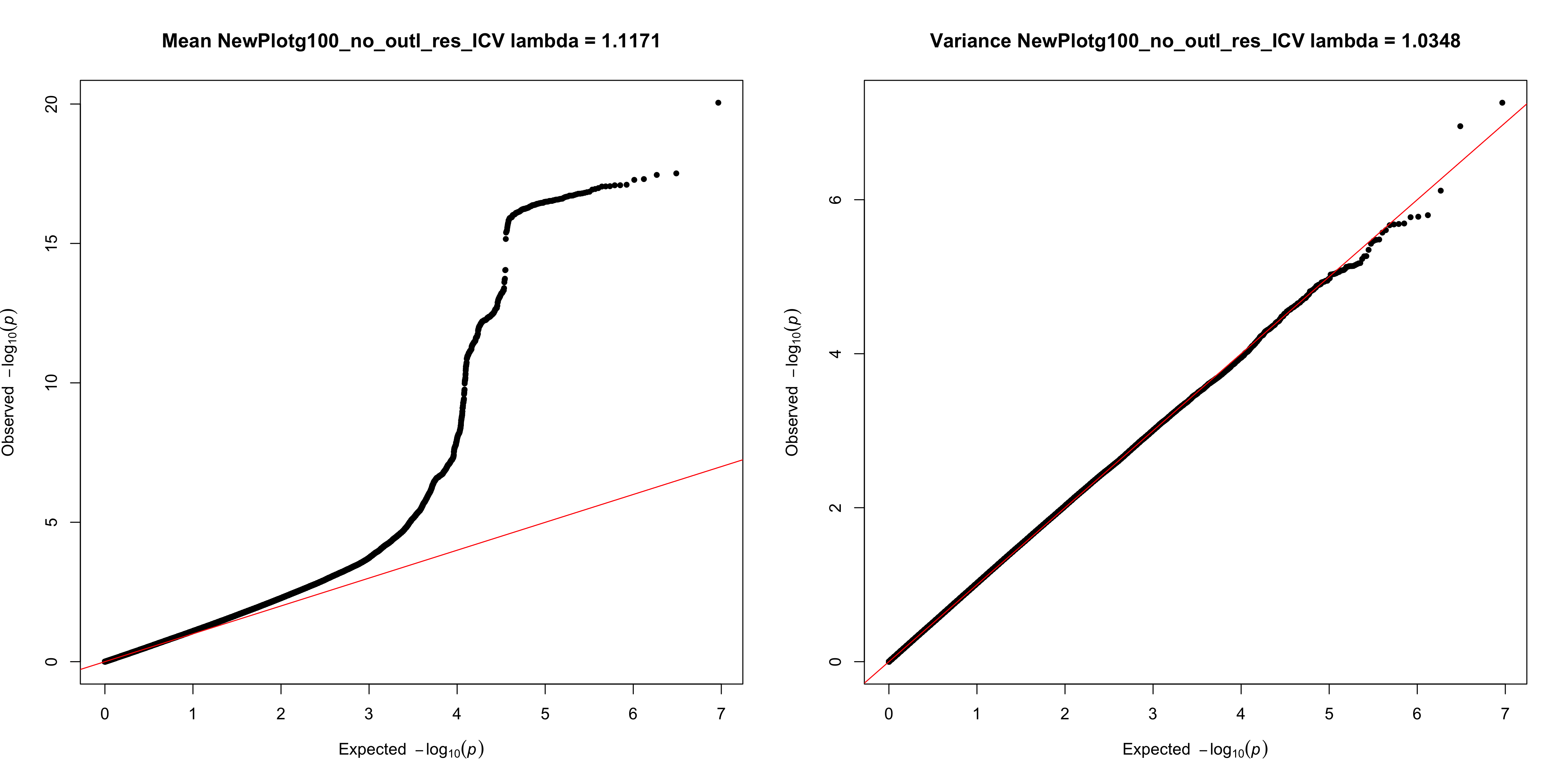
**Supplementary Figure S3i.** Manhattan and QQ plots for unadjusted mean- and variance-GWAS results, before genomic inflation correction (ICV)

**Supplementary Figure S4.** Hexagonal scatter plots of correlation between p-values from mean- and variance-models.

The plot shows -log10(p) values from mean- and variance-models, and color scales represent log_10_(count) of number of points on each bin. Red dashed lines indicate genome-wide significance thresholds (*p*=5x10^-8^) on each axis.

|  | **Mean age (s.d.)** | **Female** | **Male** | **ADHD** | **BD** | **HC** | **MDD** | **Prodr.** | **Subthreshold_**  **ADHD** | **SZ** | **SZ-SIB** | **SZBDmix** |
| --- | --- | --- | --- | --- | --- | --- | --- | --- | --- | --- | --- | --- |
| BETULA | 62.71 (13.31) | 173 | 159 | 0 | 0 | 332 | 0 | 0 | 0 | 0 | 0 | 0 |
| DEMGEN_STROKEMRI | 44.88 (22.07) | 24 | 16 | 0 | 0 | 40 | 0 | 0 | 0 | 0 | 0 | 0 |
| HUBIN | 42.07 (8.11) | 54 | 121 | 0 | 0 | 95 | 0 | 0 | 0 | 80 | 0 | 0 |
| HUNT | 58.87 (4.21) | 419 | 364 | 0 | 0 | 783 | 0 | 0 | 0 | 0 | 0 | 0 |
| NCNG | 52.49 (17.15) | 246 | 120 | 0 | 0 | 366 | 0 | 0 | 0 | 0 | 0 | 0 |
| NIMAGE | 17.68 (3.56) | 93 | 131 | 100 | 0 | 90 | 0 | 0 | 34 | 0 | 0 | 0 |
| PNC | 15.36 (3.61) | 253 | 261 | 0 | 0 | 514 | 0 | 0 | 0 | 0 | 0 | 0 |
| STROKEGE750 | 57.58 (15.40) | 69 | 48 | 0 | 0 | 117 | 0 | 0 | 0 | 0 | 0 | 0 |
| TOP15 | 33.66 (9.92) | 185 | 200 | 0 | 101 | 125 | 0 | 0 | 0 | 95 | 0 | 64 |
| TOP3T | 30.16 (9.12) | 166 | 219 | 0 | 42 | 246 | 0 | 15 | 0 | 58 | 0 | 24 |
| TOP3TGE750 | 32.14 (10.48) | 124 | 131 | 0 | 31 | 164 | 0 | 4 | 0 | 40 | 0 | 16 |
| UBA | 22.47 (3.33) | 668 | 358 | 0 | 0 | 1026 | 0 | 0 | 0 | 0 | 0 | 0 |
| UKBB | 55.67 (7.47) | 10710 | 9902 | 0 | 0 | 20612 | 0 | 0 | 0 | 0 | 0 | 0 |
| UNIBA | 28.42 (8.33) | 157 | 204 | 0 | 0 | 270 | 2 | 0 | 0 | 72 | 17 | 0 |

**Supplementary Table S2.** Distribution of sex and neuropsychiatric diagnoses across samples

Abbreviations: ADHD=attention-deficit/hyperactivity disorder, BD=bipolar disorder, HC=healthy controls, MCI=mild cognitive impairment, MDD=major depressive disorder, Prodr.=prodromal psychosis, Subtrheshold ADHD=subthreshold ADHD, SZ=schizophrenia, SZ-SIB=SZ siblings.

**REFERENCES**

Alfaro-Almagro, F., Jenkinson, M., Bangerter, N.K., Andersson, J.L.R., Griffanti, L., Douaud, G., Sotiropoulos, S.N., Jbabdi, S., Hernandez-Fernandez, M., Vallee, E.*, et al.* (2018). Image processing and Quality Control for the first 10,000 brain imaging datasets from UK Biobank. Neuroimage 166, 400-424.

Brandt, C.L., Kaufmann, T., Agartz, I., Hugdahl, K., Jensen, J., Ueland, T., Haatveit, B., Skatun, K.C., Doan, N.T., Melle, I.*, et al.* (2015). Cognitive Effort and Schizophrenia Modulate Large-Scale Functional Brain Connectivity. Schizophr Bull 41, 1360-1369.

Dørum, E.S., Alnæs, D., Kaufmann, T., Richard, G., Lund, M.J., Tønnesen, S., Sneve, M.H., Mathiesen, N.C., Rustan, Ø., Gjertsen, Ø.*, et al.* (2016). Age-related differences in brain network activation and co-activation during multiple object tracking. Brain Behav 6, e00533.

Espeseth, T., Christoforou, A., Lundervold, A.J., Steen, V.M., Le Hellard, S., and Reinvang, I. (2012). Imaging and cognitive genetics: the Norwegian Cognitive NeuroGenetics sample. Twin Res Hum Genet 15, 442-452.

Haukvik, U.K., Schaer, M., Nesvåg, R., McNeil, T., Hartberg, C.B., Jönsson, E.G., Eliez, S., and Agartz, I. (2012). Cortical folding in Broca's area relates to obstetric complications in schizophrenia patients and healthy controls. Psychol Med 42, 1329-1337.

Heck, A., Fastenrath, M., Ackermann, S., Auschra, B., Bickel, H., Coynel, D., Gschwind, L., Jessen, F., Kaduszkiewicz, H., Maier, W.*, et al.* (2014). Converging genetic and functional brain imaging evidence links neuronal excitability to working memory, psychiatric disease, and brain activity. Neuron 81, 1203-1213.

Honningsvåg, L.M., Linde, M., Håberg, A., Stovner, L.J., and Hagen, K. (2012). Does health differ between participants and non-participants in the MRI-HUNT study, a population based neuroimaging study? The Nord-Trøndelag health studies 1984-2009. BMC Med Imaging 12, 23.

Håberg, A.K., Hammer, T.A., Kvistad, K.A., Rydland, J., Müller, T.B., Eikenes, L., Gårseth, M., and Stovner, L.J. (2016). Incidental Intracranial Findings and Their Clinical Impact; The HUNT MRI Study in a General Population of 1006 Participants between 50-66 Years. PLoS One 11, e0151080.

Kaufmann, T., Alnæs, D., Doan, N.T., Brandt, C.L., Andreassen, O.A., and Westlye, L.T. (2017). Delayed stabilization and individualization in connectome development are related to psychiatric disorders. Nat Neurosci 20, 513-515.

Kaufmann, T., Skåtun, K.C., Alnæs, D., Doan, N.T., Duff, E.P., Tønnesen, S., Roussos, E., Ueland, T., Aminoff, S.R., Lagerberg, T.V.*, et al.* (2015). Disintegration of Sensorimotor Brain Networks in Schizophrenia. Schizophr Bull 41, 1326-1335.

Nilsson, L.-G., Adolfsson, R., Bäckman, L., de Frias, C.M., Molander, B., and Nyberg, L. (2004). Betula: A Prospective Cohort Study on Memory, Health and Aging. Aging, Neuropsychology, and Cognition 11, 134-148.

Pergola, G., Trizio, S., Di Carlo, P., Taurisano, P., Mancini, M., Amoroso, N., Nettis, M.A., Andriola, I., Caforio, G., Popolizio, T.*, et al.* (2017). Grey matter volume patterns in thalamic nuclei are associated with familial risk for schizophrenia. Schizophr Res 180, 13-20.

Satterthwaite, T.D., Connolly, J.J., Ruparel, K., Calkins, M.E., Jackson, C., Elliott, M.A., Roalf, D.R., Ryan Hopsona, K.P., Behr, M., Qiu, H.*, et al.* (2016). The Philadelphia Neurodevelopmental Cohort: A publicly available resource for the study of normal and abnormal brain development in youth. Neuroimage 124, 1115-1119.

Satterthwaite, T.D., Elliott, M.A., Ruparel, K., Loughead, J., Prabhakaran, K., Calkins, M.E., Hopson, R., Jackson, C., Keefe, J., Riley, M.*, et al.* (2014). Neuroimaging of the Philadelphia neurodevelopmental cohort. Neuroimage 86, 544-553.

Skåtun, K.C., Kaufmann, T., Tønnesen, S., Biele, G., Melle, I., Agartz, I., Alnæs, D., Andreassen, O.A., and Westlye, L.T. (2016). Global brain connectivity alterations in patients with schizophrenia and bipolar spectrum disorders. J Psychiatry Neurosci 41, 331-341.

von Rhein, D., Mennes, M., van Ewijk, H., Groenman, A.P., Zwiers, M.P., Oosterlaan, J., Heslenfeld, D., Franke, B., Hoekstra, P.J., Faraone, S.V.*, et al.* (2015). The NeuroIMAGE study: a prospective phenotypic, cognitive, genetic and MRI study in children with attention-deficit/hyperactivity disorder. Design and descriptives. Eur Child Adolesc Psychiatry 24, 265-281.
